# A long-lived seabird buffers the cost of phenological mismatch by modulating stress response

**DOI:** 10.1101/2025.07.10.663151

**Authors:** Flávia A. Nitta Fernandes, Gaël Bardon, Josephine R. Paris, Robin Cristofari, Joan Ferrer Obiol, Lorena Ancona, Samuele Greco, Marco Gerdol, Benoit Vallas, Pierre Carette, Céline Le Bohec, Emiliano Trucchi

## Abstract

Climate change-driven environmental variability can cause species to shift their phenology, leading to an increase in phenological mismatches. Whether the fitness cost of such mismatches can be mitigated by plasticity remains poorly understood, especially in the context of wild populations. In this study, we quantified responses to phenological mismatch using capture-mark-recapture, morphological, and transcriptomic data from king penguins hatching during and after the seasonal peak of food availability (*i.e.,* born early and late in the breeding season, respectively). Late-born chicks showed reduced survival to fledging and fledged smaller and in slightly poorer body condition than chicks born during the seasonal peak in food availability. Yet post-fledging return rates did not differ between late- and early-born chicks, suggesting that early-life stress was effectively mitigated among individuals that survived to fledging, with no detectable carryover effects in the first years post-fledging. Genes involved in oxidative stress response, homeostasis, and nutrient-sensing were differentially expressed between early- and late-born chicks, consistent with stress mitigation and homeostasis maintenance at the expense of growth in late-born chicks. A co-expression gene cluster whose primary expression pattern was associated with hatching phenology also contained central genes involved in stress mitigation. Together, our findings highlight molecular mechanisms that buffer the fitness costs of phenological mismatch, which may allow species to tolerate mismatch to some extent. Despite higher winter mortality of individuals born in mismatch, the plastic response observed in survivors hints towards molecular adaptations that could allow species resilience under increasing climate-driven environmental unpredictability.

## 1 Introduction

Rapid ongoing climate changes are associated with increasing frequency of extreme and unpredictable environmental conditions (IPCC 2023). Accelerated and intensified rates of environmental change can lead to reductions in individual’s fitness when new conditions deviate substantially from their physiological optimum (Jørgensen et al., 2022; Pörtner & Farrell, 2008), and culminate in population declines due to increased mortality (de la Fuente et al., 2025; Rao et al., 2023). Even when exposure to stressful environments does not directly result in direct mortality, such stress can impact life-long individual fitness (Crino & Breuner, 2015; Marak et al., 2003; Watson et al., 2017). Fitness impacts can be both direct, such as when thermal optima are overcome (Corregidor-Castro et al., 2023; Fangue et al., 2006; Welbergen et al., 2008), and/or indirect, such as through phenological mismatches (Li et al., 2025; Vitasse et al., 2021).

Phenological mismatches refer to the loss of synchrony in the timing of life history events between interacting species, such as prey and predator, and are expected to reduce consumer fitness (the match-mismatch hypothesis; Cushing, 1974, 1975). Moreover, according to Cushing’s classic match-mismatch hypothesis, two assumptions must be met: I) fitness is maximized when energetically demanding activities (*e.g.,* growing, breeding, migrating) occurs at the seasonal peak of resources and reduced when deviating from this peak (*i.e.*, an “in-match” baseline should exist); and II) consumers should be specialised in the seasonal resource. Although climate-driven phenological mismatches are widely documented in nature (*e.g.,* Doiron et al., 2015; Plard et al., 2014; Post & Forchhammer, 2008), only few studies meet the aforementioned assumptions (Kharouba & Wolkovich, 2023; Youngflesh et al., 2023), making it unclear to which extent mismatches actually affect species persistence.

Furthermore, the evolution of mismatch buffering strategies (*i.e.,* mechanisms that reduce the costs of phenological mismatch) may render species more resilient to climate-driven phenological shifts than previously assumed (Weir & Phillimore, 2024). Such strategies include mechanisms that reduce asynchrony between consumer and its resource (Bartomeus et al., 2013), bet-hedging strategies that render mismatches less detrimental under unpredictable environments (Schindler et al., 2015), and mechanisms that reduce the costs of mismatch in the focal species (Reed et al., 2013). Regarding the latter, in which mismatch is not avoided, but tolerated, the fitness costs of mismatch can be mitigated through ecological trade-offs (*e.g.,* lower competition when breeding late in density-dependent Great tits’ populations; Reed et al., 2013), or through physiological plasticity, such as resistance to starvation under the nutritional restrictions caused by being born out of the peak of resources (Weir & Phillimore, 2024). However, empirical evidence of phenological mismatch tolerance is still scarce, as well as the mechanisms behind mismatch mitigation.

Here, we investigated the effects of phenological mismatch and the associated molecular responses in a natural environment using the King penguin (*Aptenodytes patagonicus*) as a study system. This sub-Antarctic predator uniquely feeds on myctophid fish during the Austral summer (Cherel et al., 1993), when these fish are both the most abundant and occur within the diving range of adult king penguins (*i.e.,* 50-100 metres deep, from December to February; Charrassin & Bost, 2001; Kozlov et al., 1991). As myctophid abundance drops from May until late August, adults have to forage farther away from the colony, closer to the Antarctic pack ice (Bost et al., 2004), relying on less caloric sources of prey, such as squids (Cherel et al., 1993).

In combination with the strong resource seasonality, at every breeding event, king penguins’ hatch a single chick, with the investment in reproduction and parental care lasting up to 14 months (Descamps et al., 2002). This extended reproductive cycle results in a delayed breeding start for parents that successfully fledged a chick the previous year (Figure 1A, and Figure S1 for different years). As such, late-breeding parents lay their eggs around 1.5 months after the rest of the breeders, producing offspring at the end of the productivity peak of myctophid fish, from February on (Kozlov et al., 1991). Consequently, at each breeding event, offspring are produced in both match and mismatch with optimal food resource availability, exposing chicks within the same colony to distinct environmental conditions (Olsson, 1996; Weimerskirch et al., 1992). In addition to hatching when the peak food resource is declining, late-born chicks have less time to grow before the onset of harsh winter conditions, during which they are left alone at the colony and must fast for several months, while parents undertake longer trips to reach foraging areas (Descamps et al., 2002; Weimerskirch et al., 1992). Mismatch with food availability results in higher first-winter mortality in late-born chicks compared to early-born chicks (Figure 1C), mainly due to starvation and increased predation (Descamps et al., 2002; Weimerskirch et al., 1992).

**Figure 1.**
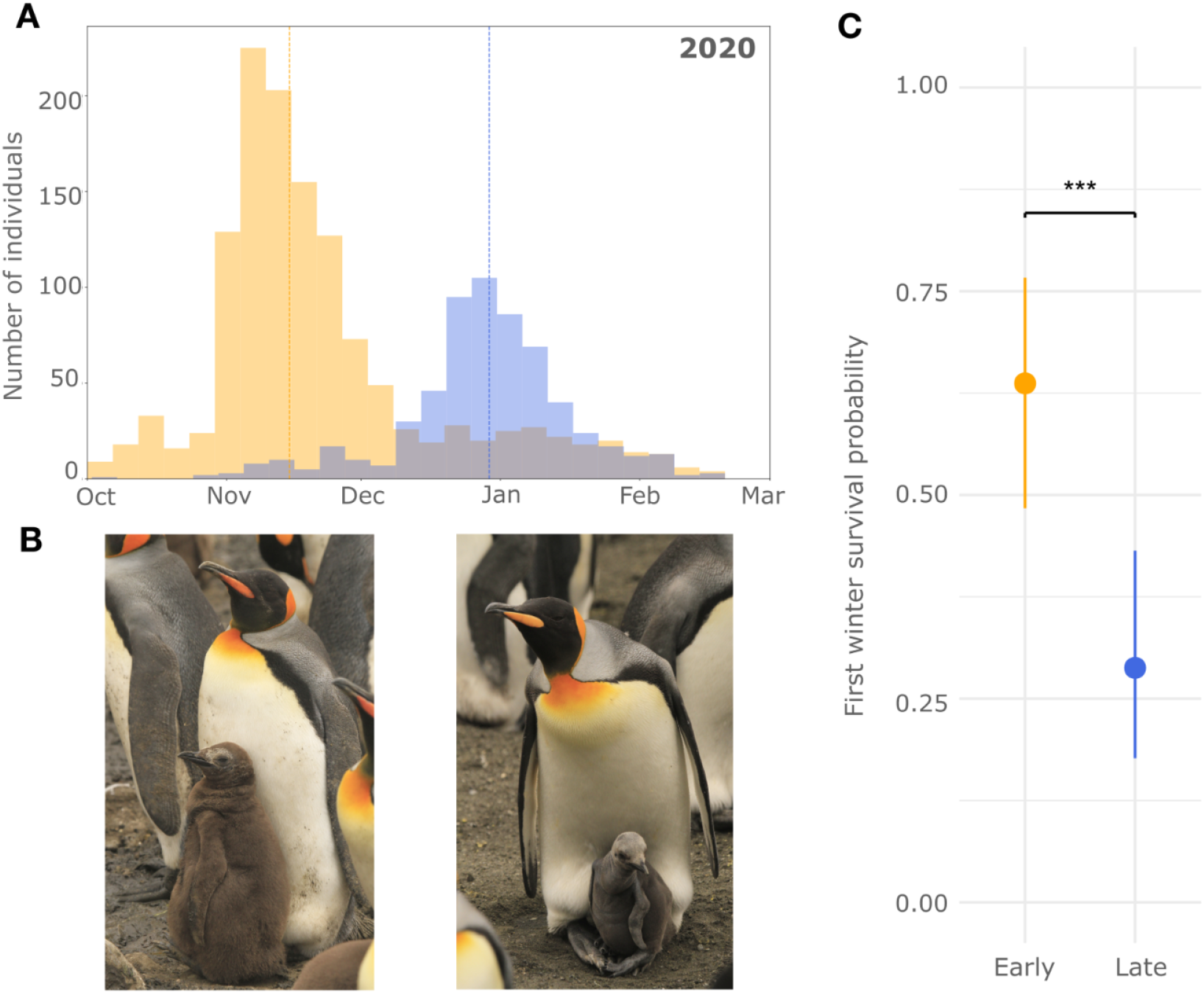
First-year survival probability linked to breeding phenology in the King penguin. **A)** Histogram of the King penguin phenology for early (orange) and late (blue) breeding individuals (defined as not successful and successful individual breeders in the previous year, respectively) in 2020, when transcriptome samples were collected (see Figure S1 for all years between 2012 and 2023). Breeding dates were defined as the onset of the individual’s first annual stay in the colony above 10 days after moulting, measured through RFID detection patterns (see Material and Methods for details). Vertical lines represent the median breeding date for each of the two groups; **B)** Early-(left) and late-born chicks (right) with a parent in the study colony (*La Baie du Marin,* Crozet archipelago). Photo taken in February 2022, when early-born chicks are thermally independent and late-born chicks remain mostly in the parent’s brooding pouch; **C)** Winter chick survival probability of early-(orange) and late-born chicks (blue). Points represent model-estimated marginal means; bars show 95% confidence intervals obtained from the mixed-effects model, with year included as a random effect.

Stressful conditions due to mismatched phenology appear to leave physiological and molecular signatures early on in development, as late-born chicks show higher levels of stress hormones and reactive oxygen species associated with shorter telomeres in the first weeks after hatching (Stier et al., 2014). Yet it remains unknown whether late-born chicks that survive the first winter can “catch-up” with early-born conspecifics in body condition at fledging and early-life survival, thereby buffering the stress caused by mismatch. To answer these questions, we harnessed long-term morphological and life-history data from 3386 individuals spanning 13 years (*i.e.,* 2010 to 2022) to assess first-winter survival of early- and late-born chicks, and to test for carryover effects of mismatch on body size and condition at fledging. We also measured early- and late-born individuals’ return rate to the breeding colony, as a proxy for survival during the first five years after fledging. Moreover, to uncover candidate gene pathways underlying potential mechanisms of mismatch buffering, we generated blood transcriptomes from 20 three-weeks-old early- and late-born chicks which successfully survived the first winter, and detected differentially expressed and co-expressed genes associated with hatching phenology.

## 2 Material and Methods

### 2.1 Long term monitoring data and analyses

Our study was conducted in the King penguin colony of *La Baie du Marin* (BDM), on Possession Island, Crozet Archipelago (46°24′27″S 51°45′27″E), in the Southern Indian Ocean. From 2010 to 2022, *ca.* 200 early-born (January) and *ca.* 200 late-born (February) chicks were captured and marked annually in the BDM sub-colony (‘Antavia’), a naturally enclosed zone with four unique passageways used by penguins to exit and enter the breeding area (Gendner et al., 2005). Passageways are equipped with underground systems of paired antennas, which capture and store the entry and exit movements of penguins equipped with Radio Frequency Identification (RFID) tags (for detailed information about this system see Bardon et al., 2023; Gendner et al., 2005). Breeding dates were defined as the beginning of stereotypical patterns of courtship and incubation in RFID detections at the individual level, corresponding to the first long annual stay at the colony (*i.e.,* > 10 days) after moulting. For more details on the estimation of breeding dates, see methods described in Bardon et al. (2023).

A total of 3386 early- and late-born chicks were sampled *ca.* 3 weeks after hatching during the brooding stage (*i.e.*, when chicks are kept in the brooding pouch of parents, being alternately guarded by one of their parents for a month). Captures occurred within a time window of 7 to 10 days, starting at the end of January for early-born chicks and at the end of February for late-born chicks. Chicks were temporarily marked with a small external plastic pin (Fishtag, *Floytag*), stamped with a unique number for individual recognition after the winter. Chicks that survived the winter were then recaptured approximately 2 weeks before fledging (*i.e.*, at the end of their first moult, at around 11 months old, between November and January) and were marked with passive RFID-tags, without any other external mark (Fishtag was removed). Winter survival was estimated from the time chicks were externally pre-marked, three weeks post-hatching, until the fledging period at the end of January.

In order to estimate carryover effects of hatching phenology on fledging phenotype, 1181 chicks, which survived the winter (out of the 3,386 that had been pre-marked), were measured for flipper length (+/- 1 mm), beak length (+/- 1 mm), and body mass (kg) at the recapture moment before fledging. For this analysis, four years of data were removed from the dataset: 2010, 2011, 2012, 2014, due to low winter survival of late-born individuals (N range from 0 to 1 chick in those years). We estimated individual body size by establishing a Structural Size Index (SSI) based on the first component of a Principal Component Analysis (PCA) of flipper and beak lengths (see Supplementary Text for details). Body condition (BC) was used as a proxy for energy storage, and was estimated from the residuals of an Ordinary Least Squares (OLS) regression of body mass on structural size (Supplementary Text). Finally, for these 1181 fledglings, we defined post-fledging return rate as the probability of being detected in the colony at least once during the five breeding seasons after fledging (starting approximately one year after departure to sea, from 2011 to 2024).

We used Generalised Linear Mixed Models (GLMMs) to test for effects of hatching timing (early-vs late-born) on fledgling size, body condition, first-winter survival rate, and post-fledging return rate. Gaussian error distributions were used for size and body condition, and binomial distributions for the two survival metrics. Hatching year was included in all models as a random effect to account for interannual variability. Model estimates, standard deviation (SD), and the significance (*P-value* < 0.05) of the explanatory variables were derived from Type-II Anova. All statistics were computed using the R v4.0.3 statistical environment (R Development Core Team, 2022).

### 2.2 Blood sampling, RNA extraction and sequencing

A specific sampling of early- and late-born chicks was conducted during the breeding season of 2020 (see Table S1 for specific dates). A total of 69 early- and 72 late-born chicks were captured and sampled. Of these, 39 early- and 10 late-born chicks survived until fledging. Chicks were sampled for blood at the brachial vein using a 25-gauge needle and a microcapillary tube. We directly transferred three to nine drops of blood into a 1.5 mL microcentrifuge tube prepared with aliquots of PAXgene® Blood RNA Solution, following the manufacturer’s recommended ratio of 2.76 blood:solution. Freezing procedures followed the manufacturer’s instructions, with final freezing at −80°C until processing in the laboratory. All manipulations were approved by the French Ethics Committee (APAFIS#4897-2015110911016428) and the French Polar Environmental Committee (TAAF permit #2019-115 & 2019-129) and conducted in accordance with their guidelines.

Total RNA was extracted from the whole blood samples using the PAXgene® Blood RNA kit, following the manufacturer’s protocol. RNA purity and concentration was assessed using a Nanodrop 2000 (Thermo Fisher Scientific, CA) and a Qubit 4.0 fluorometer (Thermo Fisher Scientific, CA), respectively. RNA integrity (RIN) was evaluated by UV transilluminator and Agilent 2100 Bioanalyzer (Agilent technologies, CA). All samples had a RIN score > 6. A total of 20 samples were selected for sequencing, 10 early-born individuals (randomly selected from the 39 survivors) and all 10 late-born individuals that survived until fledging. Libraries were prepared using the QuantSeq 3’mRNA-Seq Library Prep Kit FWD V1 (Lexogen, Vienna, Austria), a sequencing strategy that targets the 3’-end of RNA sequences, allowing the direct correspondence between read counts and transcripts per gene (Moll et al., 2014). Sequencing was performed on an Illumina NovaSeq 6000 platform in a single-end mode, targeting over 5 million reads of 75 base pairs (bp) per sample. Details about raw sequencing data and accession IDs can be found in Table S1.

### 2.3 Trimming and mapping reads to the reference genome

Raw reads were trimmed and mapped following the suggested guidelines in the Lexogen’s QuantSeq 3’ mRNA-Seq Kit and integrated Data Analysis Pipeline on Bluebee® platform. Quality of raw and cleaned reads was checked using FastQC (Andrews, 2010). Reads were mapped to the King penguin chromosome-level reference genome (GCA_965638725; Paris et al., 2025) using STAR v2.7.11b (Dobin et al., 2013), and indexed using SAMtools v1.21 (Li et al., 2009). Annotation features were counted using V1 of the species’ protein-coding annotation (Paris et al., 2025) with HTSeq v2.0.3 (Putri et al., 2022), with options *-t exon* and *-m union*. Finally, HTSeq counts for each individual were merged into a single table of raw gene counts (18,086 genes) for downstream analyses.

### 2.4 Gene counts filtering and principal component analysis

Genes aligning to the sex chromosomes (Z and W chromosomes in birds) and the highly overrepresented haemoglobin genes (HBA and HBM) were removed from the gene counts table, in order to remove major sex bias effects and dominant gene counts throughout the data, respectively. This filtered counts table was used for all downstream analyses. We proceeded by performing a principal component analysis (PCA) of the RNA expression data using broadSeq v1.0 (Das Roy, 2025). Counts were transformed using the variance stabilizing transformation (VST) and the top 500 most variable genes in expression level across all samples were selected using *prcompTidy*. Loadings of all 500 genes for the 20 principal components (PCs, one for each individual) were extracted (Table S2). PC loadings indicate to which degree each gene contributed to the variation in that PC.

### 2.5 Normalisation and differential gene expression analysis

Differential gene expression (DGE) analysis was performed between the two phenological groups of chicks using DESeq2 (Love et al., 2014). Genes with low read counts were removed by selecting only those with at least one count in at least three individuals. We observed that more than 50% of the variation in the patterns of gene expression was not due to the phenological group (Figure S2A). To investigate the likely subtle signatures of phenology in gene expression, we computed normalised gene counts with RUVSeq v1.38.0 (Risso et al., 2014). RUVSeq removes possible batch effects and unwanted variation (common in samples from wild individuals; Krishnan et al., 2020), by estimating a latent factor of unwanted variation (*W*) in the expression data, that can be added to the DGE analysis design. This latent factor is calculated from a set of control genes, which, in our case, corresponded to an empirical set of 7,019 non-significant DGEs outputted from a first DGE analysis performed prior to RUVg normalization, with the design formula: “*∼ SEX + PHENOLOGICAL_GROUP*”. Subsequently, we used the first latent factor estimated by RUVSeq, “*W_1*”, which captured the strongest dimension of unwanted variation among samples, including it to the final DGE design: “*∼ W_1 + PHENOLOGICAL_GROUP*”.

Visualisation of differentially expressed genes (DEGs) between early- and late-born chicks was done using EnhancedVolcano v1.22 (Blighe, 2018) and ComplexHeatmap v2.20 (Gu et al., 2016). Gene clustering and heatmap visualisation was performed on normalized counts for all DEGs transformed into *Z-scores*. Gene clustering was performed using the euclidean distance and complete linkage methods, corresponding to *dist* and *hclust* functions, respectively, on the *Z-score* transformed counts.

### 2.6 Co-expression module analysis

In order to identify clusters of highly correlated genes whose main expression profile was associated with late-hatching phenology, we used the weighted network construction approach of WGCNA v1.73 (Langfelder & Horvath, 2008). WGCNA allows the construction of gene networks and the identification of modules of genes (*i.e.*, clusters of genes with correlated gene expression) that can be associated with external traits of interest. We followed the standard procedure in WGCNA of defining a soft-thresholding power, a parameter used to weight gene-gene correlations, such that strong correlations are amplified and weak correlations are downweighted (Zhang & Horvath, 2005). Briefly, this step allows highlighting more robust co-expression relationships. The RUVSeq normalised counts table was used as an input. We selected a power of 16 for network construction, corresponding to the value for which the scale-free topology model fit (R^2^) was higher than 0.85 (Horvath & Dong, 2008).

A gene co-expression network was constructed for both early- and late-born individuals using the *blockwiseModules* function from WGCNA (Langfelder & Horvath, 2012). We set the *blockwiseModules* parameters to *networkType = “signed”*, in order to exclusively detect positive correlations amongst genes, avoiding the confounding influence of negative correlations (Mason et al., 2009; van Dam et al., 2018), *minModuleSize = 30* to filter out modules with less than 30 genes, *corType= “bicor”* (*i.e.*, bidweight midcorrelation, median-based measure of similarity between samples), *mergeCutHeight = 0.25* (cut height for module merging in the dendrogram), and kept all other parameters as default.

We then calculated the correlation between the main expression profiles of co-expression modules and hatching phenology, as well as sex, the latter as a control for modules that could be biassed towards one sex. A Pearson correlation (*cor* function) was used with *P-values* determined using a Student’s asymptotic test (*corPvalueStudent* function) between traits and module eigengenes (MEs), which corresponds to each module’s first principal component (PC1) of expression profile (Langfelder & Horvath, 2008). We considered a module to be positively correlated with a trait (early-born vs. late-born, or male vs. female) if *cor* > 0.5 and *P-value* < 0.05.

Finally, we identified hub genes (*i.e.*, central genes, highly connected in the given module) in modules correlated with hatching phenology. Hub genes were defined as genes whose intramodular connectivity (kIM) was above the 90% quantile of the kIM distribution for that module, following the same procedure used in Paris et al. (2024). The higher the kIM value (closer to 1), the more connected a gene is within its module.

### 2.7 Gene ontology enrichment

In order to interpret the potential biological function of modules of interest and DEGs, we performed gene ontology (GO) enrichment analyses using clusterProfiler v4.12.6 (Xu et al., 2024) *enricher* function. We used the chicken (*Gallus gallus*) gene sets and GO terms as reference, which were downloaded from Ensembl using biomart v2.56 (Durinck et al., 2009), setting *minGSSize* to 5 and *maxGSSize* to 500, and an FDR-corrected *P-value* ≤ 0.1. Biological process GO terms were summarised into broader GO processes using the function *goSlim()* from the R package GSEABase v1.66.0 (Morgan et al., 2024). GO slims simplify the overview of functional biological processes, by summarising GO terms (child GO terms) into higher level GO terms (parent GO terms). Relationships between parent GOs, child GOs and genes that enriched the child GOs were plotted using *ggalluvial* v0.12.5 (Brunson, 2020).

## 3 Results

### 3.1 Phenological mismatch affects early-life body size, but not survival after fledging

Harnessing 13 years of capture-mark-recapture data, we found that early-born chicks had a substantially higher survival probability during the first winter (0.64, 95% CI: 0.60–0.67) compared with late-born chicks (0.29, 95% CI: 0.26–0.31; GLMM – Binomial: *P-value* < 0.001, N = 3386; Figure 1C; Table S3). Additionally, late-born chicks that survived the first winter showed carryover effects on the winter developmental growth, measured as structural body size index (SSI) and body condition (BC) at fledging (see Supplementary Text for details). SSI was markedly higher in early-born chicks compared to late-born chicks, after adjusting for interannual variation (GLMM – Gaussian: *P-value* < 0.001, N = 1181; Figure 2A; Table S3). Hatching date also had an effect on fledging BC, although statistical significance was weaker (GLMM – Gaussian: *P-value* = 0.042, N = 1181; Figure 2B; Table S3). On average, early-born chicks had a slightly higher BC compared to late-hatched chicks, after adjusting for the year (Table S3).

**Figure 2.**
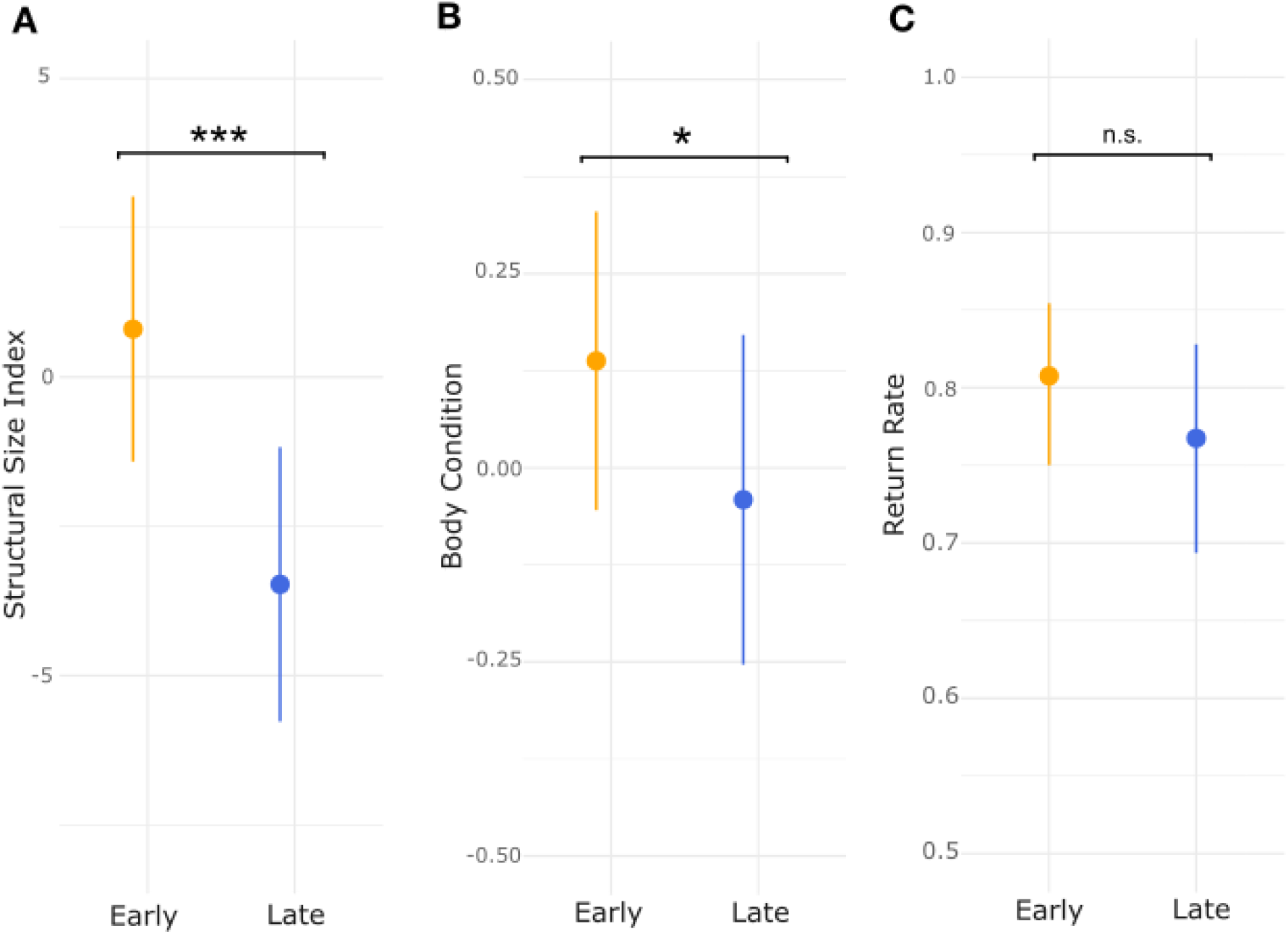
Carryover effects of mismatch on fledging morphology and five-year post-fledging survival. Estimated marginal means from three generalized linear mixed models (GLMMs) assessing the influence of hatching date (early vs. late) on fledging and post-fledging traits in king penguins. Points represent model-estimated marginal means; bars show 95% confidence intervals obtained from the GLMM, with year included as a random effect. Panels A) and B) show effects on **A)** fledging structural size and **B)** body condition (Gaussian GLMMs). **C)** Probability of return to the breeding colony from the year of fledging until September 2024, where 0 was assigned to individuals that never returned after fledging and 1 was assigned to individuals that returned at least once within five years after fledging. n.s. denotes a non-significant effect.

Despite the significant differences in body size and body condition at fledging, return rates of early- and late-born individuals across five years after fledging did not significantly differ (GLMM – Binomial: *P-value* = 0.113, N = 1181; Table S3), in spite of a trend of early-born individuals having higher return rates (Figure 2C). To further explore how the observed differences in BC affect post-fledging survival, we tested a model including BC as the sole predictor of return probability. The model predicted only a 0.3% difference in return probability between groups (early-born: 0.797; late-born: 0.794). These results suggest that, although fledging BC differed significantly between early- and late-born chicks, the effect size was too small to meaningfully influence post-fledging survival prospects.

### 3.2 Higher stress response, homeostasis maintenance, and cell differentiation in late-born chicks

In order to identify molecular mechanisms that could allow equal post-fledging survival despite being born in mismatch, we generated whole blood transcriptomic data for 10 early- and 10 late-born chicks sampled three weeks after hatching (QuantSeq 3’mRNA-Seq, see Table S1). After read trimming, quality filtering and mapping, we analysed the expression data of 16,904 genes, excluding genes in sex chromosomes and overrepresented haemoglobin genes (see Material and Methods for details). Principal component analysis of the 500 most variable genes revealed segregation between early- and late-born chicks along the third principal component (PC3), which explained 7.68% of the total gene expression variance (Figure 3A, Figure S2B). PC1 and PC2 did not separate the biological replicates according to any measured biological traits (Figure S2A, PC1=38.47%, PC2=18.22% of the variance explained). As our data harbours gene expression information from the different cell types present in the blood (*i.e.,* nucleated erythrocytes, thrombocytes and different white blood cells; Maxwell et al., 2024), cell type could be driving the variation explained by PC1 and PC2 in our data set. Additionally, the transcript heterogeneity that is characteristic of wild individuals reared in non-controlled conditions (Krishnan et al., 2020) could have overwhelmed the first two PCs. Yet, blood remains one of the least invasive tissues that can be collected while allowing quantification of stress responses in birds (Watson et al., 2017).

**Figure 3.**
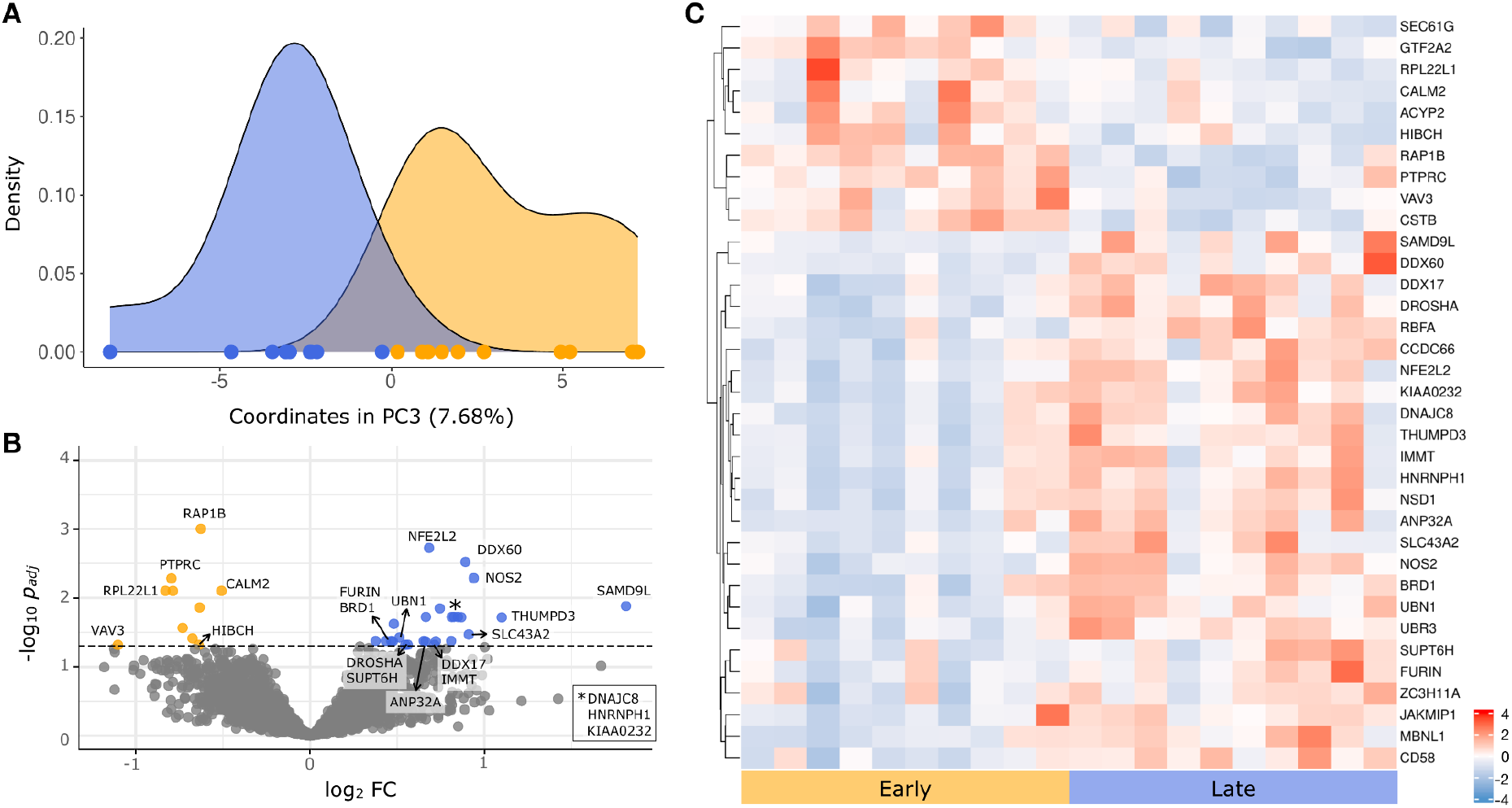
Differentially expressed genes between early- and late-born king penguin chicks. **A)** Density plot of PC3 gene expression amongst the samples (early-born chicks in orange, late-born chicks in blue) using the 500 most variable genes (see Figure S2B for sample distribution in PC3-PC4 coordinates); **B)** Volcano plot of the 5,355 genes used in the differential gene expression analysis. Differentially-expressed genes (DEGs) between the two phenological groups, where orange shows the significant underexpressed (10) genes and blue shows the significant overexpressed (25) genes in late-born chicks; **C)** Heatmap showing *Z-scores* of RUVSeq normalised read counts per gene for the 35 DEGs between early- and late-born chicks. Gene clustering was performed using euclidean distance and complete linkage methods after *Z-score* transformation of the expression data. Sample clustering was based on hatching phenology, not on gene expression profiles.

Using a set of 5,355 missing data-filtered genes (Figure S3, see Material and Methods for details), we detected 35 differentially expressed genes (DEGs) between early- and late-born chicks, with 25 genes being over-expressed and 10 genes being under-expressed in late-born chicks (Figure 3B,C, Table S4). In accordance with the variance explained by the third PC (7.68%), the log_2_ fold changes of expression between the two phenological groups were generally low, ranging from 0.37 to 1.81 (Table S4).

RAP1B was the most significant DEG (log2FC=−0.63, *p-adj*=9.9E-4; Table S4), being underexpressed in late-born chicks. This gene belongs to the evolutionarily conserved Rap1-GTPase protein family and is regulated by stimuli such as growth factors and cytokines (*i.e.,* regulators of inflammation), showing a diversity of downstream functions (*e.g.,* platelet aggregation, cell-cell contact modulation; reviewed by Frische & Zwartkruis, 2010). RAP1B is also a suppressor of the nutrient signalling pathway mTORC1 complex during amino acid starvation (Mutvei et al., 2020), a pathway that regulates nutrient signaling, cell growth, and metabolism (Panwar et al., 2023).

Late-born chicks also underexpressed genes involved in blood cell development, such as PTPRC and RPL22L1, the latter of which is required for the emergence of hematopoietic stem cells (Zhang et al., 2013). The next most significantly underexpressed gene in late-born chicks was CALM2 (calmodulin 2), which is involved in the maintenance of erythrocyte homeostasis in chickens (Alves-Ferreira et al., 1999).

Among the most significantly overexpressed genes in late-born chicks, we detected two regulators of stress response: the antioxidant protein transcription factor NFE2L2 (log2FC=−0.68, *p-adj*=1.89E-3), functionally related to oxidative stress mitigation, neurodegeneration, and ageing (Bell et al., 2016; Goodfellow et al., 2020); and the nitric oxide synthase NOS2 (log2FC=−0.94, *p-adj*=5.16E-3), which encodes an enzyme involved in the production of nitric oxide, a reactive nitrogen species that plays a key role in regulating energetic metabolism (Sansbury & Hill, 2014). As the underexpressed RAP1B gene, NOS2 also regulates the mTORC1 pathway. However, in contrast with RAP1B, NOS2 activates the mTORC1 signalling pathway through autophagy inhibition (Sarkar et al., 2011). The detected antagonistic expression patterns of NOS2 and RAP1B could indicate the activation of the mTORC1 pathway in late-born chicks during the sampling period. Additionally, DDX60 (log2FC=−0.94, *p-adj*=5.16E-3), a DEAD box protein related to antiviral immune response and cellular homeostasis (Tapescu & Cherry, 2024), was the second most highly significantly overexpressed gene in late-born chicks.

In regards to the potential functional roles of differentially expressed genes, we detected microRNA (miRNA) metabolic process as the top enriched biological process (fold enrichment = 134.9, *P-adjusted* = 0.016) among a total of 103 GOs enriched by the 35 DEGs (Table S5). This term included two overexpressed genes in late-born chicks: a DEAD box protein, DDX17, and DROSHA, both of which are known to control miRNA biogenesis, a process crucial for metabolic homeostasis (Fukuda et al., 2007). DDX17 knockdown from human hematopoietic stem and progenitor cells is also known to result in higher exposure to DNA damage (Jagdhane et al., 2019), suggesting this gene’s potential function in the maintenance of cell homeostasis under stress.

A total of 30 broader biological processes (*i.e.,* GO slims) could be summarised from the 103 GO terms (Figure S4, Table S5). The largest GO slims included immune system response, signaling, anatomical structure development, and cell differentiation (Figure S4). Biological functions within these four main GO slims showed highly complementary functions, as immune system response terms were also involved in blood cell differentiation and signaling (Figure 4). Most biological functions were related to blood cell differentiation, mainly through the activity of BRD1 (overexpressed in late chicks, regulating the differentiation of erythrocytes) and PTPRC (underexpressed in late chicks, regulating the differentiation of other nucleated blood cells).

**Figure 4.**
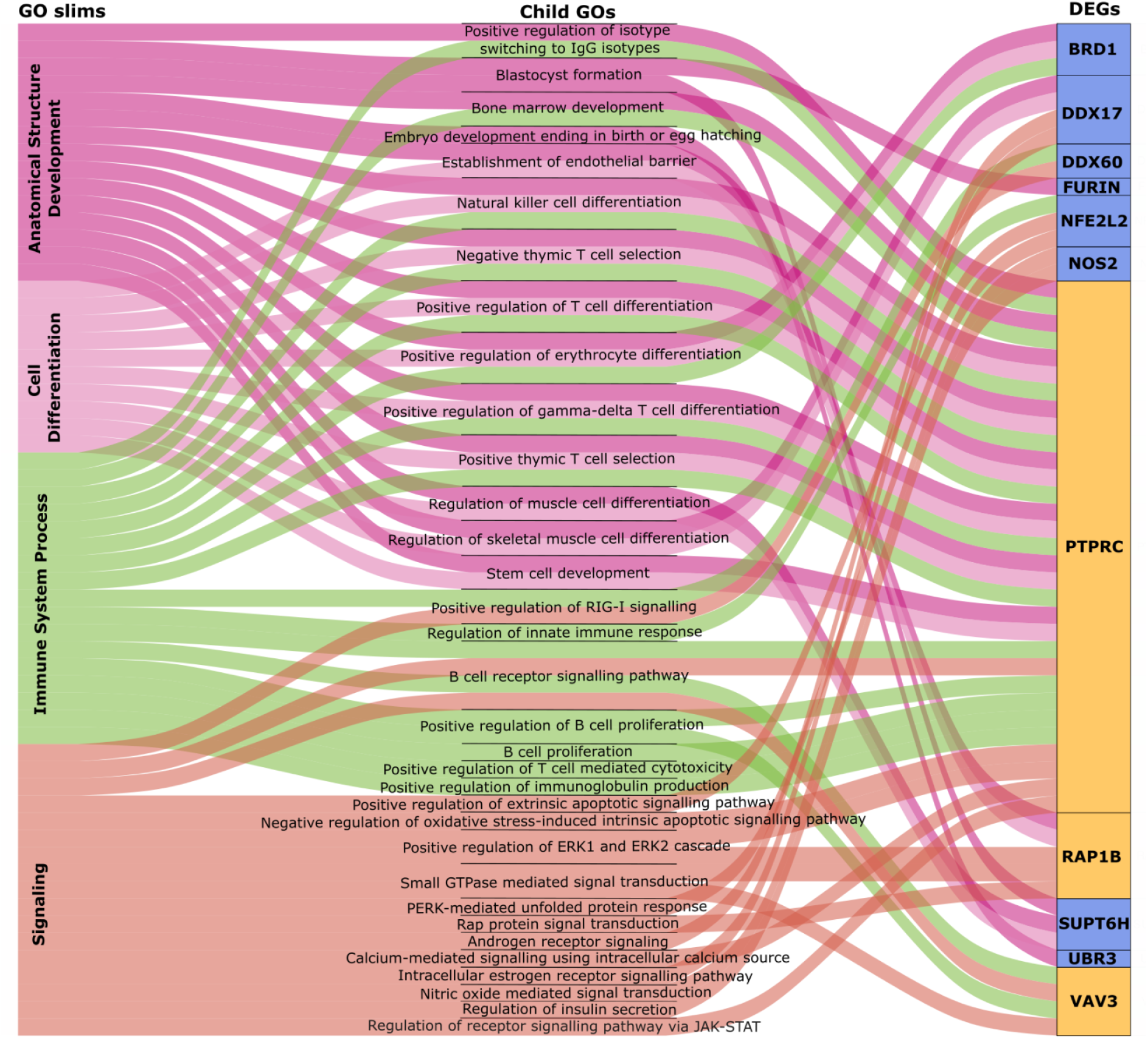
Main molecular functions affected by mismatched hatching phenology. Sankey plot of the top four GO slims and their child GO terms enriched by the differentially expressed genes. Alluvials are coloured by the four GO slims and the DEGs enriching GO terms are coloured by overexpression in the different phenological groups (orange: overexpressed genes in early-born chicks; blue: overexpressed genes in late-born chicks).

Other biological processes not intrinsically related to the blood transcriptome and immunity were also detected, such as regulation of muscle cell differentiation, insulin secretion, and oxidative stress response pathways, all of which included overexpressed genes in late-born individuals (Figure 4, Table S5).

### 3.3 Coordinated activity of stress-mitigation genes in late-born chicks

To further investigate the potential functional implication of DEGs through their association with the expression of other genes, we generated a gene correlation network of early- and late-born chicks expression data (N = 5,355 genes). We detected eight clusters of genes with highly correlated expression profiles, *i.e.,* co-expression modules, from the gene correlation network (Table S6). Of the eight co-expression modules detected, seven contained genes previously identified as differentially expressed between early- and late-born chicks: modules 1, 3, and 7 contained overexpressed genes in late-born chicks, while modules 2, 4, and 6 contained overexpressed genes in the early-born ones (Figure 5A, Table S6). Module 7 (N = 407 genes) contained 14 out of 25 overexpressed genes in late-born chicks, including the aforementioned antioxidants NOS2 and NFE2L2, as well as homeostasis regulators DDX60 and DROSHA (Table S7). This was the only module to exhibit a significant difference in gene expression profiles between early- and late-born chicks, based on the module eigengene (ME), a measure of the primary axis of gene expression variation in that module (Figure 5B, Table S8). Accordingly, this module’s ME was positively correlated with late-hatching phenology (Figure 5C, *cor* = 0.59, *P-value* = 0.006). Other modules showed positive (modules 1 and 3) and negative (module 5) correlation trends between ME and late-hatching phenology, but these were not statistically significant (Table S8). Module 5 was also correlated with sex (*P-value* = 0.02, Figure S6A, Table S6), but ME was not significantly different between females and males (*P-value* = 0.34, Figure S6B, Table S8; see Supplementary Text section on Sex variation for further details). We thus proceeded by exploring the potential functional role of module 7, positively associated with late-hatching phenology.

**Figure 5.**
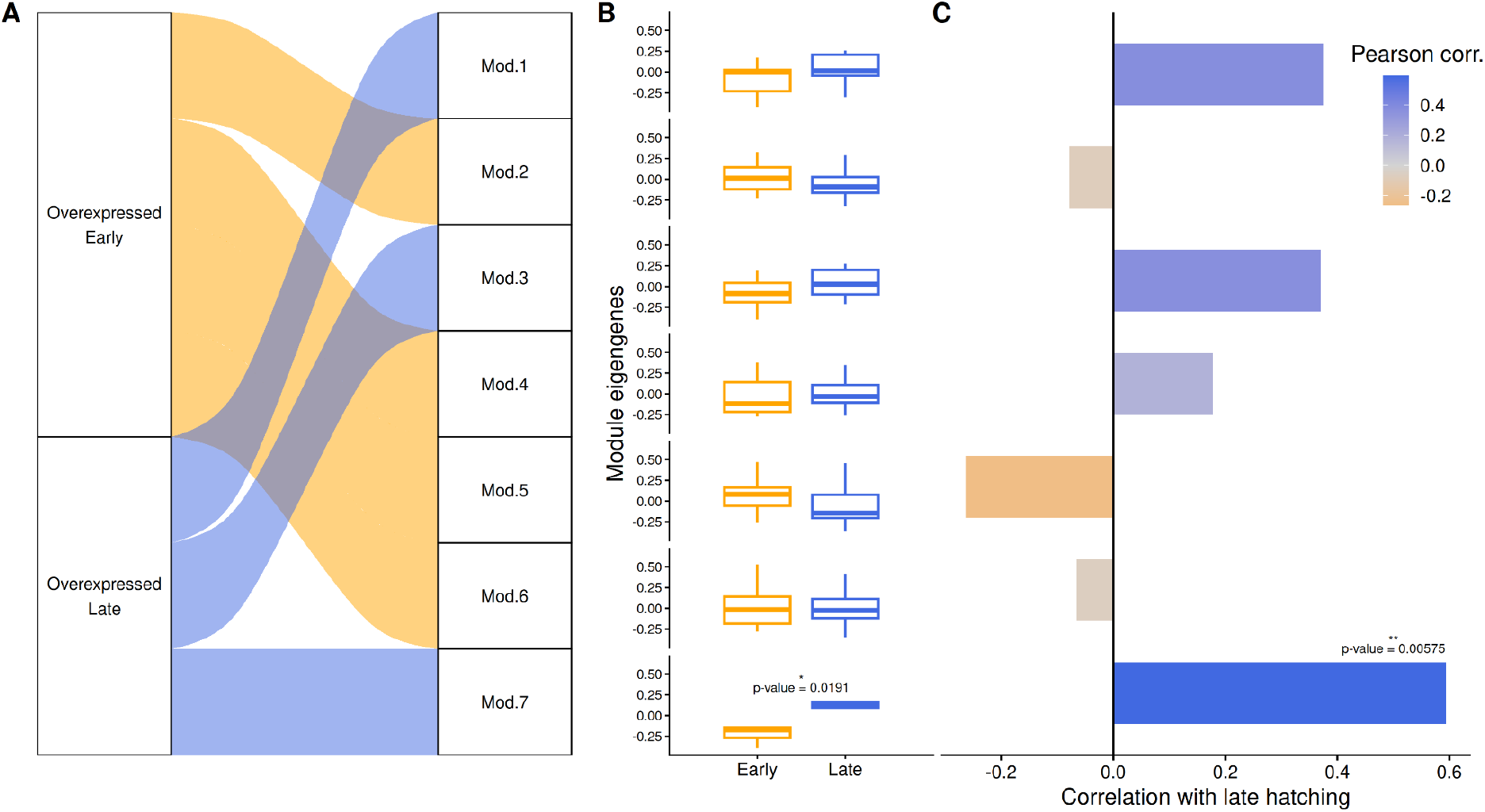
Correlation of co-expression modules with phenology. **A)** Sankey plot showing the distribution of differentially expressed genes in each co-expression module (overexpressed genes in late-born chicks: blue; overexpressed genes in early-born chicks: orange); **B)** Boxplots summarize the distribution of module eigengene values (MEs) within early- and late-born chicks. Module eigengenes represent the overall expression profile of each co-expression module, summarizing the combined expression patterns of genes within that module. P-value indicates Kruskal-Wallis tests rank sum test results for differences between mean MEs between the two phenological groups; Detailed statistics in Table S8; **C)** Pearson correlation between MEs and late-hatching phenology (darker blue colors represent stronger correlations with late-hatching); Detailed statistics in Table S6. The y-axes in panels (B) and (C) follow the module order shown in panel (A).

Gene ontology enrichment of the 407 genes in the module associated with late-hatching phenology resulted in ten GO terms (Table S9). Protein refolding was the top biological process, containing five heat shock protein genes, known for their key role in the maintenance of homeostasis in organisms under stress responses (Hartl et al., 2011). Cellular response to glucose starvation and endoplasmic reticulum unfolded protein response were the subsequent top GO terms, which included an overlapping set of genes related to stress response: the heat shock protein HSPA5, which was also enriched in the protein refolding term; the translation initiation factor EIF2AK3; the antioxidant protein transcription factor NFE2L2 (overexpressed in late-born chicks); and the mTORC1-regulated transcription factor ATF4, which regulates cellular response to metabolic stress. Although not differentially expressed between early- and late-born individuals, the ATF4 transcription factor is a direct target of the mTORC1 metabolic pathway (Liu & Sabatini, 2020), which has regulators that are over- and underexpressed in the different groups of chicks (RAP1B, NOS2), as previously discussed.

Alongside gene ontology enrichment analyses, we identified highly connected (“hub”) genes within the co-expression module associated with late-hatching phenology, as hub genes are associated with the modules’ primary functional roles (Langfelder & Horvath, 2008). Among the 40 hub genes identified (intra-module connectivity, kIM_Hub genes_ > 0.67, Table S10), the most central hub gene was ERC1 (kIM = 1), also known as ELKS, which functions as a key regulator of a conserved DNA damage signaling pathway (Wu et al., 2010). The splicing factor 3b subunit 1 (SF2B1, kIM = 0,92) and the heat shock protein DNAJC8 (kIM=0.90), were the following most central hub genes in this module, with DNAJC8 also being overexpressed in late-born chicks (Table S7). Seven other overexpressed genes in late-hatchlings were detected as hub genes in this module (Table S7). Based on previous studies, these genes are functionally related to homeostasis; IMMT (Liu et al., 2024), HNRNPH1 (Brownmiller & Caplen, 2023); immune response; ANP32A; (Baker et al., 2018), HNRNPH1; (Tong et al., 2021); and stress: DNAJC8 (Fatima et al., 2026), NFE2L2 (Ryoo & Kwak, 2018), THUMPD3 (Tan et al., 2022).

## 4 Discussion

The extent to which species can mitigate the negative effects of phenological mismatches remains understudied (Weir & Phillimore, 2024), limiting our ability to assess the fitness costs of the increasing phenological shifts observed in recent decades (Kharouba & Wolkovich, 2020, 2023). Here, we demonstrate that king penguins born in mismatch with the peak of food resources buffer the effects of mismatch, at least partially, as they had similar post-fledging survival prospects compared to individuals born during the peak of resources, despite higher mortality during the first winter. Mismatch buffering mechanisms appear to involve the overexpression and co-regulation of genes involved in stress response, immunity, and homeostasis, in late-born chicks. The challenge of maintaining homeostasis while facing stressful mismatched conditions during development was, however, accompanied by the disruption of blood cell development and growth pathways, underexpressed in late-born chicks. High stress levels can restrict growth due to the direct impact of stressors in growth pathways (Sadoul & Vijayan, 2016; Sävendahl, 2012), as well as through the increased energy allocated for homeostatic maintenance (Faught et al., 2016). Accordingly, late-born individuals also fledged smaller and in slightly worse body conditions than individuals born during the resource peak.

Although body condition at fledging is a predictor of survival in many bird species (Blums et al., 2005; Conroy et al., 1989; Hill et al., 2003; Pace & Afton, 1999), including king penguins (Saraux et al., 2011), our findings do not directly contradict this expectation, as differences in body condition were barely significant and effect sizes not sufficiently strong to affect return rates. Smaller body size, on the other hand, is not expected to have a negative effect on survival, although the relationship between body size and survival varies among bird species (Wilcox et al., 2024). Producing smaller offspring is less energetically costly for breeders under resource-limited conditions (Williams, 1966), a pattern that has previously been suggested for king penguins in years of harsher environmental conditions (Bordier et al., 2014). Likewise, producing smaller chicks may be a less costly strategy for late-breeding parents, not only due to the challenge of providing food when myctophid resources are scarcer (Kozlov et al., 1991), but also due to the accumulated reproductive costs from successfully fledging a chick in the previous year (Le Bohec et al., 2007).

Furthermore, smaller body sizes could be a consequence of energetic trade-offs in late-born chicks, especially during the shorter window of time these chicks have from hatching to the winter fasting (Descamps et al., 2002; Weimerskirch et al., 1992). Late-born chicks seem to allocate resources to maintenance tasks, such as homeostasis, oxidative stress repair, and immune response, instead of investing in growth, a commonly observed trade-off under limited resources (Pontzer & McGrosky, 2022). Despite immune response expression patterns being unclear, with key genes for blood cell development (*i.e.,* PTPRC, RPL22L1, and CALM2) being underexpressed in late-born chicks, and others being overexpressed (*i.e.,* DDX60 and BRD1), oxidative stress mitigation was the main observed response in this group of chicks.

The detection of overexpressed genes related to stress (notably NFE2L2, NOS2) indicates higher oxidative stress in late-born chicks, alongside mechanisms to mitigate this stress, including the overexpression of the DNA damage repair gene DDX17. Additionally, the detection of DNA damage regulator ERC1 as a central regulator of heat shock protein coding genes, antioxidants, as well as regulators of homeostasis and immune response in the co-expression module 7, associated with late hatching, reinforces the hypothesis of a coordinated stress mitigation response in late-born chicks a few weeks after hatching. Regulatory changes in Hsp60 heat shock proteins have been previously detected under physiological stress caused by poor food intake in an experimental study in juvenile white ibis (Herring et al., 2011). Although we lack information about differential caloric intake between early- and late-born chicks during the first year of life, we detected DEGs related to nutrient transport and metabolic stress (SLC43A2; Zhang et al., 2023), as well as selenium deficiency (PTPRC; Lingamgunta et al., 2023), while module 7 had a significantly enriched GO term related to cellular response to glucose starvation (Table S8).

In line with this, the overexpression of NOS2 (activator of mTORC1 and regulator of autophagy under amino acid starvation) and the underexpression of RAP1B (suppressor of mTORC1 during amino acid starvation) indicate a potential activation of the mTORC1 nutrient pathway in late-born chicks due to nutrient privation. mTORC1 activation is usually related with cell growth demands and here, is likely involved in a feedback loop to maintain cells in a nutritionally sparse environment (Liu & Sabatini, 2020). Chronic mTOR pathway activation is linked with accelerated ageing (Liu & Sabatini, 2020), but the increased activity of mTOR in early-life stages does not necessarily mean it is chronically activated throughout life, nor that late-born individuals are subjected to accelerated ageing.

Whether the observed early-life stress experienced by late-born individuals leads to carryover effects on survival and reproduction throughout their lifespan is currently not testable with our data. The first cohort of hatching chicks have been followed since 2010 in a species where the first breeding attempt occurs around 5 years of age (Le Bohec, 2007), and the average lifespan in the wild is *ca.* 20 years (Gauthier–Clerc et al., 2004). Furthermore, in addition to the mTOR regulators, three overexpressed genes in late-born chicks were also related with cellular senescence and longevity: SAMD9L, UBN1, and NFE2L2. The regulator of cell proliferation SAMD9L has convergently evolved in birds and bats, being potentially linked with increased lifespan despite high metabolic rates in flying taxa (Matsuda & Makino, 2024). Ubinuclein 1, UBN1, is a component of a chromatin-remodeling pathway needed for proper neural differentiation in mice (Xiong et al., 2018), but also a key cellular senescence regulator (Banumathy et al., 2009). Due to the non-senescent nature of the three-week-old chicks from our study, we hypothesise that the overexpression of UBN1 is mostly due to intense cellular differentiation. Finally, NFE2L2, one of the hub genes in co-expression module 7 and also related to oxidative stress mitigation, is related to neurodegeneration and ageing (Bell et al., 2016; Goodfellow et al., 2020). Despite the association between hatching group and expression of these genes, their functional implications remain to be explored.

Here we demonstrate that king penguins harbour plastic molecular adaptations that buffer environmental mismatches through the interaction of growth and stress mitigation pathways. These buffering mechanisms seemingly facilitate post-fledging survival, albeit at the expense of reduced body size at fledging, suggesting a trade-off between growth and the maintenance of physiological homeostasis. Mismatch buffering could also be a consequence of stronger viability selection on late-born chicks, indicated by higher winter mortality rates, which would select for individuals who are able to grow under lower nutrient intake and higher stress levels (Stier et al., 2014). Accordingly, adult king penguins naturally endure prolonged fasting periods during moulting and incubation (Gauthier-Clerc et al., 2002). Future studies could test whether late-born chicks that survive to adulthood exhibit greater resistance to fasting than individuals born earlier in the season. On the one hand, the observed adaptive plastic response could reflect long-term past adaptation to highly variable environmental conditions, while, on the other hand, play a key role in the species’ persistence under unpredictable future climate conditions, when phenological mismatches are expected to become more frequent.

Uncovering the molecular basis of phenological mismatch buffering is essential for understanding how species may respond to climate-driven phenological shifts. While meta-analyses have been fundamental in quantifying the prevalence and fitness consequences of phenological mismatches in nature (Kharouba & Wolkovich, 2023; Thackeray et al., 2016), our findings highlight the importance of species-specific studies to identify the physiological and molecular processes that underpin resilience to mismatch. Elevated stress levels are also experienced by individuals born late in the breeding season in other long-lived seabirds (Kitaysky et al., 1999), suggesting that the plastic responses identified in king penguins may represent a more general mechanism for buffering the effects of phenological mismatch across species, which should be further tested.

## Acknowledgments

This study was supported by the Institut Polaire Français Paul-Emile Victor (IPEV) within the framework of the project 137-ANTAVIA-POLAROBS, by the Centre Scientifique de Monaco, and by the Centre National de la Recherche Scientifique (CNRS) through the Programme Zone Atelier de Recherches sur l’Environnement Antarctique et Terres Australes Subantarctique (ZATA). This study is part of and supported by the long-term Studies in Ecology and Evolution (SEE-Life) programme of CNRS. FANF was supported by the PhD scholarship of DiSVA/UNIVPM under a joint agreement with the University of Strasbourg, and the Research Council of Finland grant #354649 to Robin Cristofari. We are deeply grateful to all the wintering and summering members of project IPEV 137 and all the other colleagues and students within the team, who participated in the long-term monitoring as part of the project IPEV 137 since 1998. All members of the MIBE team at the IPHC in Strasbourg are thanked for their technical expertise and support. We also sincerely thank the IPEV logistics teams for their important and continued support in the field. This study was approved by the French Polar Environmental Committee and permits handling animals and access breeding sites were delivered by the Terres Australes et Antarctiques Françaises (TAAF). Computational analyses were performed at DISVA-HPC and CSC – IT Center for Science, Finland.

## Data Accessibility Statement

Raw sequencing reads are deposited in the SRA database within BioProject ID PRJNA1288185, and Bio-Sample numbers: SAMN49836463-SAMN49836482.

## Benefit-Sharing Statement

The authors declare that the Nagoya Protocol is not applicable to this research and that there are no benefits to report.

## Author contributions

Conceptualization: FANF, GB, CLB, ET. Methodology: FANF, GB, CLB, ET. Data collection: FANF, GB, BV, PC, RC, CLB, ET. Data analysis: FANF, GB. Supervision: CLB, ET. Writing—original draft: FANF, GB, JRP, JFO, CLB, ET. Writing—edited manuscript: FANF, GB, RC, CLB, ET. All authors contributed critically to the drafts and gave final approval for publication.

## Ethics

Health, safety, security and other risks for participating researchers were assessed and managed by the institution (Institut Polaire Français Paul-Emile Victor, IPEV) that provided logistic support at the research stations involved. Ethics questions concerning local and regional researchers, partners, or governments do not apply, due to Antarctica being uninhabited.

## References

Alves-Ferreira, M., Scofano, H. M., & Ferreira-Pereira, A. (1999). Ca(2+)-ATPase from chicken (Gallus domesticus) erythrocyte plasma membrane: effects of calmodulin and taurine on the Ca(2+)-dependent ATPase activity and Ca2+ uptake. Comparative Biochemistry and Physiology. Part B, Biochemistry & Molecular Biology, 122(3), 269–276.

Andrews, S. (2010). FastQC A Quality Control tool for High Throughput Sequence Data. https://www.bioinformatics.babraham.ac.uk/projects/fastqc/

Baker, S. F., Ledwith, M. P., & Mehle, A. (2018). Differential splicing of ANP32A in birds alters its ability to stimulate RNA synthesis by restricted influenza polymerase. Cell Reports, 24(10), 2581–2588.e4.

Banumathy, G., Somaiah, N., Zhang, R., Tang, Y., Hoffmann, J., Andrake, M., Ceulemans, H., Schultz, D., Marmorstein, R., & Adams, P. D. (2009). Human UBN1 is an ortholog of yeast Hpc2p and has an essential role in the HIRA/ASF1a chromatin-remodeling pathway in senescent cells. Molecular and Cellular Biology, 29(3), 758–770.

Bardon, G., Cristofari, R., Winterl, A., Barracho, T., Benoiste, M., Ceresa, C., Chatelain, N., Courtecuisse, J., Fernandes, F. A. N., Gauthier-Clerc, M., Gendner, J.-P., Handrich, Y., Houstin, A., Krellenstein, A., Lecomte, N., Salmon, C.-E., Trucchi, E., Vallas, B., Wong, E. M., … Le Bohec, C. (2023). RFIDeep: Unfolding the potential of deep learning for radio-frequency identification. Methods in Ecology and Evolution / British Ecological Society. 10.1111/2041-210x.14187

Bartomeus, I., Park, M. G., Gibbs, J., Danforth, B. N., Lakso, A. N., & Winfree, R. (2013). Biodiversity ensures plant-pollinator phenological synchrony against climate change. Ecology Letters, 16(11), 1331–1338.

Bell, C. G., Xia, Y., Yuan, W., Gao, F., Ward, K., Roos, L., Mangino, M., Hysi, P. G., Bell, J., Wang, J., & Spector, T. D. (2016). Novel regional age-associated DNA methylation changes within human common disease-associated loci. Genome Biology, 17(1), 193.

Blighe, K. (2018). EnhancedVolcano: Publication-ready volcano plots with enhanced colouring and labeling. Github. https://github.com/kevinblighe/EnhancedVolcano

Blums, P., Nichols, J. D., Hines, J. E., Lindberg, M. S., & Mednis, A. (2005). Individual quality, survival variation and patterns of phenotypic selection on body condition and timing of nesting in birds. Oecologia, 143(3), 365–376.

Bordier, C., Saraux, C., Viblanc, V. A., Gachot-Neveu, H., Beaugey, M., Le Maho, Y., & Le Bohec, C. (2014). Inter-Annual Variability of Fledgling Sex Ratio in King Penguins. PloS One, 9(12), e114052.

Bost, C. A., Charrassin, J. B., Clerquin, Y., Ropert-Coudert, Y., & Le Maho, Y. (2004). Exploitation of distant marginal ice zones by king penguins during winter. Marine Ecology Progress Series, 283, 293–297.

Brownmiller, T., & Caplen, N. J. (2023). The HNRNPF/H RNA binding proteins and disease. Wiley Interdisciplinary Reviews. RNA, 14(5), e1788.

Brunson, J. C. (2020). ggalluvial: Layered Grammar for Alluvial Plots. Journal of Open Source Software, 5(49), 2017.

Charrassin, J. B., & Bost, C. A. (2001). Utilisation of the oceanic habitat by king penguins over the annual cycle. Marine Ecology Progress Series, 221, 285–298.

Cherel, Y., Verdon, C., & Ridoux, V. (1993). Seasonal importance of oceanic myctophids in king penguin diet at Crozet Islands. Polar Biology, 13(5). 10.1007/bf00238362

Conroy, M. J., Costanzo, G. R., & Stotts, D. B. (1989). Winter survival of female American black ducks on the Atlantic coast. The Journal of Wildlife Management, 53(1), 99.

Corregidor-Castro, A., Morinay, J., McKinlay, S. E., Ramellini, S., Assandri, G., Bazzi, G., Glavaschi, A., De Capua, E. L., Grapputo, A., Romano, A., Morganti, M., Cecere, J. G., Pilastro, A., & Rubolini, D. (2023). Experimental nest cooling reveals dramatic effects of heatwaves on reproduction in a Mediterranean bird of prey. Global Change Biology, 29(19), 5552–5567.

Crino, O. L., & Breuner, C. W. (2015). Developmental stress: evidence for positive phenotypic and fitness effects in birds. Journal of Ornithology / DO-G, 156(1), 389–398.

Cushing, D. H. (1974). The natural regulation of fish populations. In: F.R. Harden Jones (Ed.). Sea Fisheries Research, 399–412.

Cushing, D. H. (1975). Marine ecology and fisheries. Cambridge University Press.

Das Roy, R. (2025). broadSeq. Bioconductor. http://bioconductor.org/packages/broadSeq/

de la Fuente, A., Briscoe, N. J., Kearney, M. R., Williams, S. E., Youngentob, K. N., Marsh, K. J., Cernusak, L. A., Leahy, L., Larson, J., & Krockenberger, A. K. (2025). Climate-induced physiological stress drives Rainforest mammal population declines. Global Change Biology, 31(5), e70215.

Descamps, S., Gauthier-Clerc, M., Gendner, J. P., & Le Maho, Y. (2002). The annual cycle of unbanded king penguins Aptenodytes patagonicus on Possession Island (Crozet). 2, 87–98.

Dobin, A., Davis, C. A., Schlesinger, F., Drenkow, J., Zaleski, C., Jha, S., Batut, P., Chaisson, M., & Gingeras, T. R. (2013). STAR: ultrafast universal RNA-seq aligner. Bioinformatics, 29(1), 15–21.

Doiron, M., Gauthier, G., & Lévesque, E. (2015). Trophic mismatch and its effects on the growth of young in an Arctic herbivore. Global Change Biology, 21(12), 4364–4376.

Durinck, S., Spellman, P. T., Birney, E., & Huber, W. (2009). Mapping identifiers for the integration of genomic datasets with the R/Bioconductor package biomaRt. Nature Protocols, 4(8), 1184–1191.

Fangue, N. A., Hofmeister, M., & Schulte, P. M. (2006). Intraspecific variation in thermal tolerance and heat shock protein gene expression in common killifish, Fundulus heteroclitus. The Journal of Experimental Biology, 209(Pt 15), 2859–2872.

Fatima, S., Gupta, A., & Priya, S. (2026). Stress-specific expression of HSPA and DNAJ chaperones regulated by NRF2/HSF1 in neurodegenerative conditions. Molecular and Cellular Biochemistry, 481(4), 1661–1676.

Faught, E., Aluru, N., & Vijayan, M. M. (2016). The Molecular Stress Response. In Fish Physiology (Vol. 35, pp. 113–166). Elsevier.

Frische, E. W., & Zwartkruis, F. J. T. (2010). Rap1, a mercenary among the Ras-like GTPases. Developmental Biology, 340(1), 1–9.

Fukuda, T., Yamagata, K., Fujiyama, S., Matsumoto, T., Koshida, I., Yoshimura, K., Mihara, M., Naitou, M., Endoh, H., Nakamura, T., Akimoto, C., Yamamoto, Y., Katagiri, T., Foulds, C., Takezawa, S., Kitagawa, H., Takeyama, K.-I., O’Malley, B. W., & Kato, S. (2007). DEAD-box RNA helicase subunits of the Drosha complex are required for processing of rRNA and a subset of microRNAs. Nature Cell Biology, 9(5), 604–611.

Gauthier–Clerc, M., Gendner, J.-P., Ribic, C. A., Fraser, W. R., Woehler, E. J., Descamps, S., Gilly, C., Le Bohec, C., & Le Maho, Y. (2004). Long–term effects of flipper bands on penguins. Proceedings of the Royal Society of London. Series B: Biological Sciences, 271(suppl_6), S423–S426.

Gauthier-Clerc, M., Le Maho, Y., Gendner, J.-P., & Handrich, Y. (2002). Moulting fast and time constraint for reproduction in the king penguin. Polar Biology, 25(4), 288–295.

Gendner, J.-P., Gauthier-Clerc, M., Le Bohec, C., Descamps, S., & Le Maho, Y. (2005). A new application for transponders in studying penguins. Journal of Field Ornithology, 76(2), 138–142.

Goodfellow, M. J., Borcar, A., Proctor, J. L., Greco, T., Rosenthal, R. E., & Fiskum, G. (2020). Transcriptional activation of antioxidant gene expression by Nrf2 protects against mitochondrial dysfunction and neuronal death associated with acute and chronic neurodegeneration. Experimental Neurology, 328(113247), 113247.

Gu, Z., Eils, R., & Schlesner, M. (2016). Complex heatmaps reveal patterns and correlations in multidimensional genomic data. *Bioinformatics (Oxford*, England), 32(18), 2847–2849.

Hartl, F. U., Bracher, A., & Hayer-Hartl, M. (2011). Molecular chaperones in protein folding and proteostasis. Nature, 475(7356), 324–332.

Herring, G., Cook, M. I., Gawlik, D. E., & Call, E. M. (2011). Food availability is expressed through physiological stress indicators in nestling white ibis: a food supplementation experiment: Food availability and physiological stress in ibis. Functional Ecology, 25(3), 682–690.

Hill, M. R. J., Alisauskas, R. T., Ankney, C. D., & Leafloor, J. O. (2003). Influence of body size and condition on harvest and survival of juvenile Canada geese. The Journal of Wildlife Management, 67(3), 530.

Horvath, S., & Dong, J. (2008). Geometric interpretation of gene coexpression network analysis. PLoS Computational Biology, 4(8), e1000117.

Jagdhane, P., Garg, S., Xia, J., He, L., Misiak, D., Mortusewicz, O., Helleday, T., Müller-Tidow, C., Koehn, M., & Pabst, C. (2019). Pf230 the RNA helicase ddx17 protects acute myeloid leukemia cells against DNA damage: PF230. HemaSphere, 3(S1), 67.

Jørgensen, L. B., Ørsted, M., Malte, H., Wang, T., & Overgaard, J. (2022). Extreme escalation of heat failure rates in ectotherms with global warming. Nature, 611(7934), 93–98.

Kharouba, H. M., & Wolkovich, E. M. (2020). Disconnects between ecological theory and data in phenological mismatch research. Nature Climate Change, 10(5), 406–415.

Kharouba, H. M., & Wolkovich, E. M. (2023). Lack of evidence for the match-mismatch hypothesis across terrestrial trophic interactions. Ecology Letters, 26(6), 955–964.

Kitaysky, A. S., Wingfield, J. C., & Piatt, J. F. (1999). Dynamics of food availability, body condition and physiological stress response in breeding Black-legged Kittiwakes: Food availability and stress response in Kittiwakes. Functional Ecology, 13(5), 577–584.

Kozlov, A. N., Shust, K. V., & Zemsky, A. V. (1991). Seasonal and inter·annual variability in the distribution of Electrona carlsbergi in the southern polar front area (the area to the north of South Georgia is used as an example) (Nos. 7, 337–368). CCAMLR Sel Sci Pap. https://www.ccamlr.org/en/system/files/science_journal_papers/21-Kozlov-et-al.pdf

Krishnan, J., Persons, J. L., Peuß, R., Hassan, H., Kenzior, A., Xiong, S., Olsen, L., Maldonado, E., Kowalko, J. E., & Rohner, N. (2020). Comparative transcriptome analysis of wild and lab populations of Astyanax mexicanus uncovers differential effects of environment and morphotype on gene expression. Journal of Experimental Zoology. Part B, Molecular and Developmental Evolution, 334(7-8), 530–539.

Langfelder, P., & Horvath, S. (2008). WGCNA: an R package for weighted correlation network analysis. BMC Bioinformatics, 9, 559.

Langfelder, P., & Horvath, S. (2012). Fast R Functions for Robust Correlations and Hierarchical Clustering. Journal of Statistical Software, 46(11). 10.18637/jss.v046.i11

Le Bohec, C. (2007). Stratégies d’histoire de vie d’un oiseau longévif: Le manchot royal (Aptenodytes Patagonicus) [Université Louis Pasteur (Strasbourg) (1971-2008)]. https://theses.fr/2007STR13234

Le Bohec, C., Gauthier-Clerc, M., Grémillet, D., Pradel, R., Béchet, A., Gendner, J.-P., & Le Maho, Y. (2007). Population dynamics in a long-lived seabird: I. Impact of breeding activity on survival and breeding probability in unbanded king penguins. The Journal of Animal Ecology, 76(6), 1149–1160.

Li, D., Belitz, M., Campbell, L., & Guralnick, R. (2025). Extreme weather events have strong but different impacts on plant and insect phenology. Nature Climate Change, 15(3), 321–328.

Li, H., Handsaker, B., Wysoker, A., Fennell, T., Ruan, J., Homer, N., Marth, G., Abecasis, G., Durbin, R., & 1000 Genome Project Data Processing Subgroup. (2009). The Sequence Alignment/Map format and SAMtools. Bioinformatics, 25(16), 2078–2079.

Lingamgunta, L. K., Aloor, B. P., Dasari, S., Ramakrishnan, R., Botlagunta, M., Madikonda, A. K., Gopal, S., & Sade, A. (2023). Identification of prognostic hub genes and therapeutic targets for selenium deficiency in chicks model through transcriptome profiling. Scientific Reports, 13(1), 8695.

Liu, G. Y., & Sabatini, D. M. (2020). mTOR at the nexus of nutrition, growth, ageing and disease. Nature Reviews. Molecular Cell Biology, 21(4), 183–203.

Liu, L., Zhao, Q., Xiong, D., Li, D., Du, J., Huang, Y., Yang, Y., & Chen, R. (2024). Suppressing mitochondrial inner membrane protein (IMMT) inhibits the proliferation of breast cancer cells through mitochondrial remodeling and metabolic regulation. Scientific Reports, 14(1), 12766.

Love, M. I., Huber, W., & Anders, S. (2014). Moderated estimation of fold change and dispersion for RNA-seq data with DESeq2. Genome Biology, 15(12), 550.

Marak, H. B., Biere, A., & Van Damme, J. M. M. (2003). Fitness costs of chemical defense in Plantago lanceolata L.: effects of nutrient and competition stress. Evolution; International Journal of Organic Evolution, 57(11), 2519–2530.

Mason, M. J., Fan, G., Plath, K., Zhou, Q., & Horvath, S. (2009). Signed weighted gene co-expression network analysis of transcriptional regulation in murine embryonic stem cells. BMC Genomics, 10(1), 327.

Matsuda, Y., & Makino, T. (2024). Comparative genomics reveals convergent signals associated with the high metabolism and longevity in birds and bats. *Proceedings*. Biological Sciences, 291(2029), 20241068.

Maxwell, M., Söderlund, R., Härtle, S., & Wattrang, E. (2024). Single-cell RNA-seq mapping of chicken peripheral blood leukocytes. BMC Genomics, 25(1), 124.

Moll, P., Ante, M., Seitz, A., & Reda, T. (2014). QuantSeq 3′ mRNA sequencing for RNA quantification. Nature Methods, 11(12), i – iii.

Morgan, M., Falcon, S., & Gentleman, R. (2024). GSEABase. Bioconductor. http://bioconductor.org/packages/GSEABase/

Mutvei, A. P., Nagiec, M. J., Hamann, J. C., Kim, S. G., Vincent, C. T., & Blenis, J. (2020). Rap1-GTPases control mTORC1 activity by coordinating lysosome organization with amino acid availability. Nature Communications, 11(1), 1416.

Olsson, O. (1996). Seasonal Effects of Timing and Reproduction in the King Penguin: A Unique Breeding Cycle. Journal of Avian Biology, 27(1), 7–14.

Pace, R. M., & Afton, A. D. (1999). Direct recovery rates of lesser scaup banded in northwest Minnesota: Sources of heterogeneity. The Journal of Wildlife Management, 63(1), 389.

Panwar, V., Singh, A., Bhatt, M., Tonk, R. K., Azizov, S., Raza, A. S., Sengupta, S., Kumar, D., & Garg, M. (2023). Multifaceted role of mTOR (mammalian target of rapamycin) signaling pathway in human health and disease. Signal Transduction and Targeted Therapy, 8(1), 375.

Paris, J. R., Fernandes, F. A. N., Santos, C. A., Pointon, D.-L. B., Wood, J. M. D., Obiol, J. F., Salces-Ortiz, J., Fernández, R., Cristofari, R., Le Bohec, C., & Trucchi, E. (2025). A chromosome-level genome of the King penguin (*Aptenodytes patagonicus*): an emerging model-in-the-wild for studying evolution. In bioRxiv (p. 2025.03. 13.642884). 10.1101/2025.03.13.642884

Paris, J. R., Nitta Fernandes, F. A., Pirri, F., Greco, S., Gerdol, M., Pallavicini, A., Benoiste, M., Cornec, C., Zane, L., Haas, B., Le Bohec, C., & Trucchi, E. (2024). Gene expression shifts in Emperor penguin adaptation to the extreme Antarctic environment. *Molecular Ecology*, e17552.

Plard, F., Gaillard, J.-M., Coulson, T., Hewison, A. J. M., Delorme, D., Warnant, C., & Bonenfant, C. (2014). Mismatch between birth date and vegetation phenology slows the demography of roe deer. PLoS Biology, 12(4), e1001828.

Pontzer, H., & McGrosky, A. (2022). Balancing growth, reproduction, maintenance, and activity in evolved energy economies. Current Biology, 32(12), R709–R719.

Pörtner, H. O., & Farrell, A. P. (2008). Physiology and climate change. *Science (New York*, N.Y.), 322(5902), 690–692.

Post, E., & Forchhammer, M. C. (2008). Climate change reduces reproductive success of an Arctic herbivore through trophic mismatch. Philosophical Transactions of the Royal Society of London. Series B, Biological Sciences, 363(1501), 2369–2375.

Putri, G. H., Anders, S., Pyl, P. T., Pimanda, J. E., & Zanini, F. (2022). Analysing high-throughput sequencing data in Python with HTSeq 2.0. *Bioinformatics (Oxford*, England), 38(10), 2943–2945.

Rao, M. P., Davi, N. K., Magney, T. S., Andreu-Hayles, L., Nachin, B., Suran, B., Varuolo-Clarke, A. M., Cook, B. I., D’Arrigo, R. D., Pederson, N., Odrentsen, L., Rodríguez-Catón, M., Leland, C., Burentogtokh, J., Gardner, W. R. M., & Griffin, K. L. (2023). Approaching a thermal tipping point in the Eurasian boreal forest at its southern margin. Communications Earth & Environment, 4(1), 247.

Reed, T. E., Grøtan, V., Jenouvrier, S., Sæther, B.-E., & Visser, M. E. (2013). Population growth in a wild bird is buffered against phenological mismatch. Science (New York, N.Y.), 340(6131), 488–491.

Risso, D., Ngai, J., Speed, T. P., & Dudoit, S. (2014). Normalization of RNA-seq data using factor analysis of control genes or samples. Nature Biotechnology, 32(9), 896–902.

Ryoo, I.-G., & Kwak, M.-K. (2018). Regulatory crosstalk between the oxidative stress-related transcription factor Nfe2l2/Nrf2 and mitochondria. Toxicology and Applied Pharmacology, 359, 24–33.

Sadoul, B., & Vijayan, M. M. (2016). Stress and Growth. In Fish Physiology (Vol. 35, pp. 167–205). Elsevier.

Sansbury, B. E., & Hill, B. G. (2014). Regulation of obesity and insulin resistance by nitric oxide. Free Radical Biology & Medicine, 73, 383–399.

Saraux, C., Viblanc, V. A., Hanuise, N., Le Maho, Y., & Le Bohec, C. (2011). Effects of individual pre-fledging traits and environmental conditions on return patterns in juvenile king penguins. PloS One, 6(6), e20407.

Sarkar, S., Korolchuk, V. I., Renna, M., Imarisio, S., Fleming, A., Williams, A., Garcia-Arencibia, M., Rose, C., Luo, S., Underwood, B. R., Kroemer, G., O’Kane, C. J., & Rubinsztein, D. C. (2011). Complex inhibitory effects of nitric oxide on autophagy. Molecular Cell, 43(1), 19–32.

Sävendahl, L. (2012). The effect of acute and chronic stress on growth. Science Signaling, 5(247), t9.

Schindler, D. E., Armstrong, J. B., & Reed, T. E. (2015). The portfolio concept in ecology and evolution. Frontiers in Ecology and the Environment, 13(5), 257–263.

Stier, A., Viblanc, V. A., Massemin-Challet, S., Handrich, Y., Zahn, S., Rojas, E. R., Saraux, C., Le Vaillant, M., Prud’homme, O., Grosbellet, E., Robin, J.-P., Bize, P., & Criscuolo, F. (2014). Starting with a handicap: phenotypic differences between early- and late-born king penguin chicks and their survival correlates. Functional Ecology, 28(3), 601–611.

Tan, Y., Liu, L., Zhang, X., Xue, Y., Gao, J., Zhao, J., Chi, N., & Zhu, Y. (2022). THUMPD3-AS1 is correlated with gastric cancer and regulates cell function through miR-1252-3p and *CXCL17*. Critical Reviews in Eukaryotic Gene Expression, 32(8), 69–80.

Tapescu, I., & Cherry, S. (2024). DDX RNA helicases: key players in cellular homeostasis and innate antiviral immunity. Journal of Virology, 98(10), e0004024.

Thackeray, S. J., Henrys, P. A., Hemming, D., Bell, J. R., Botham, M. S., Burthe, S., Helaouet, P., Johns, D. G., Jones, I. D., Leech, D. I., Mackay, E. B., Massimino, D., Atkinson, S., Bacon, P. J., Brereton, T. M., Carvalho, L., Clutton-Brock, T. H., Duck, C., Edwards, M., … Wanless, S. (2016). Phenological sensitivity to climate across taxa and trophic levels. Nature, 535(7611), 241–245.

Tong, L., Chu, Z., Gao, X., Yang, M., Adam, F. E. A., Theodore, D. W. P., Liu, H., Wang, X., Xiao, S., & Yang, Z. (2021). Newcastle disease virus V protein interacts with hnRNP H1 to promote viral replication. Veterinary Microbiology, 260(109093), 109093.

van Dam, S., Võsa, U., van der Graaf, A., Franke, L., & de Magalhães, J. P. (2018). Gene co-expression analysis for functional classification and gene-disease predictions. Briefings in Bioinformatics, 19(4), 575–592.

Vitasse, Y., Ursenbacher, S., Klein, G., Bohnenstengel, T., Chittaro, Y., Delestrade, A., Monnerat, C., Rebetez, M., Rixen, C., Strebel, N., Schmidt, B. R., Wipf, S., Wohlgemuth, T., Yoccoz, N. G., & Lenoir, J. (2021). Phenological and elevational shifts of plants, animals and fungi under climate change in the European Alps. Biological Reviews of the Cambridge Philosophical Society, 96(5), 1816–1835.

Watson, H., Videvall, E., Andersson, M. N., & Isaksson, C. (2017). Transcriptome analysis of a wild bird reveals physiological responses to the urban environment. Scientific Reports, 7, 44180.

Weimerskirch, H., Stahl, J. C., & Jouventin, P. (1992). The breeding biology and population dynamics of King Penguins Aptenodytes patagonica on the Crozet Islands. The Ibis, 134(2), 107–117.

Weir, J. C., & Phillimore, A. B. (2024). Buffering and phenological mismatch: A change of perspective. Global Change Biology, 30(5), e17294.

Welbergen, J. A., Klose, S. M., Markus, N., & Eby, P. (2008). Climate change and the effects of temperature extremes on Australian flying-foxes. *Proceedings*. Biological Sciences, 275(1633), 419–425.

Wilcox, R. C., Benson, T. J., Brawn, J. D., & Tarwater, C. E. (2024). Observed declines in body size have differential effects on survival and recruitment, but no effect on population growth in tropical birds. Global Change Biology, 30(8), e17455.

Williams, G. C. (1966). Natural Selection, the Costs of Reproduction, and a Refinement of Lack’s Principle. The American Naturalist, 100(916), 687–690.

Wu, Z.-H., Wong, E. T., Shi, Y., Niu, J., Chen, Z., Miyamoto, S., & Tergaonkar, V. (2010). ATM- and NEMO-dependent ELKS ubiquitination coordinates TAK1-mediated IKK activation in response to genotoxic stress. Molecular Cell, 40(1), 75–86.

Xiong, C., Wen, Z., Yu, J., Chen, J., Liu, C.-P., Zhang, X., Chen, P., Xu, R.-M., & Li, G. (2018). UBN1/2 of HIRA complex is responsible for recognition and deposition of H3.3 at cis-regulatory elements of genes in mouse ES cells. BMC Biology, 16(1), 110.

Xu, S., Hu, E., Cai, Y., Xie, Z., Luo, X., Zhan, L., Tang, W., Wang, Q., Liu, B., Wang, R., Xie, W., Wu, T., Xie, L., & Yu, G. (2024). Using clusterProfiler to characterize multiomics data. Nature Protocols, 19(11), 3292–3320.

Youngflesh, C., Montgomery, G. A., Saracco, J. F., Miller, D. A. W., Guralnick, R. P., Hurlbert, A. H., Siegel, R. B., LaFrance, R., & Tingley, M. W. (2023). Demographic consequences of phenological asynchrony for North American songbirds. Proceedings of the National Academy of Sciences of the United States of America, 120(28), e2221961120.

Zhang, B., & Horvath, S. (2005). A general framework for weighted gene co-expression network analysis. Statistical Applications in Genetics and Molecular Biology, 4, Article17.

Zhang, C., Shafaq-Zadah, M., Pawling, J., Hesketh, G. G., Dransart, E., Pacholczyk, K., Longo, J., Gingras, A.-C., Penn, L. Z., Johannes, L., & Dennis, J. W. (2023). SLC3A2 N-glycosylation and Golgi remodeling regulate SLC7A amino acid exchangers and stress mitigation. The Journal of Biological Chemistry, 299(12), 105416.

Zhang, Y., Duc, A.-C. E., Rao, S., Sun, X.-L., Bilbee, A. N., Rhodes, M., Li, Q., Kappes, D. J., Rhodes, J., & Wiest, D. L. (2013). Control of hematopoietic stem cell emergence by antagonistic functions of ribosomal protein paralogs. Developmental Cell, 24(4), 411–425.

