## Supplementary Text for "A long-lived seabird buffers the cost of phenological mismatch by modulating stress response"

#### Individual traits: morphological measurements and body condition

Measurements of flipper length (± 1 mm), beak length (± 1 mm), and body mass (kg) were collected at fledging (N=1181 chicks). Flipper and beak lengths represent good proxies of the penguin's structural size and are known to be highly correlated (Fahlman et al. 2006). We established a Structural Size Index (SSI) using the first component of a principal component analysis of flipper and beak lengths, as previously described in Saraux and collaborators (Saraux et al. 2011). The following equation, obtained from measures collected at fledging, was used to calculate the SSI at fledging:

$$SSI \;=\; PC1 \;=\; 0.30 * Beak \;+\; 0.95 \;*\; Flipper$$

Body mass is highly variable in this species, reflecting differences in nutritional status as well as structural size(Saraux et al. 2011). In fact, an individual can have a high mass because it has a large structural size or because it is carrying metabolised energetic reserves in the form of fat or protein (Dobson 1992). When calculating the individual's energy store through its body mass, we must correct for structural body size. Therefore, we used an Ordinary Least Squares (OLS) regression residuals of body mass on structural size to provide a better reflection of the actual energy stores of the animal (Schulte-Hostedde et al. 2005; Saraux et al. 2011; Bordier et al. 2014). Individuals with positive residuals are considered to be in better body condition (BC) and have higher energy storage than individuals with negative residuals (Jakob et al. 1996; Schulte-Hostedde et al. 2001).

#### Sex variation

Despite removal of sex-chromosome-associated genes, we also identified a co-expression module correlated with sex (module 5, Figure S6). This module, composed of 774 genes, was enriched for two GO terms: actin cytoskeleton organization (17 genes, *P-adjusted* = 0.038) and actin filament bundle assembly (6 genes, *P-adjusted* = 0.060). Actin cytoskeleton activity has been previously shown to control sex steroids remodeling in human neurons (Sanchez et al. 2011) and could be related to sex differentiation. Although king penguins do not show observable sex dimorphism (Kriesell et al. 2018), further investigation is required to fully understand the still under-explored autosomal control of sex determination (Gunski et al. 2017), which was not the focus of this study. These sex-related genes could be a statistical artefact, or alternatively could imply the identification of potentially sexually-antagonistic loci (Dutoit et al. 2018) or hitherto under-explored autosomal control of sex determination (Gunski et al. 2017). Further investigation is required to fully understand the relationship of these genes with sex.

### Supplementary Figures

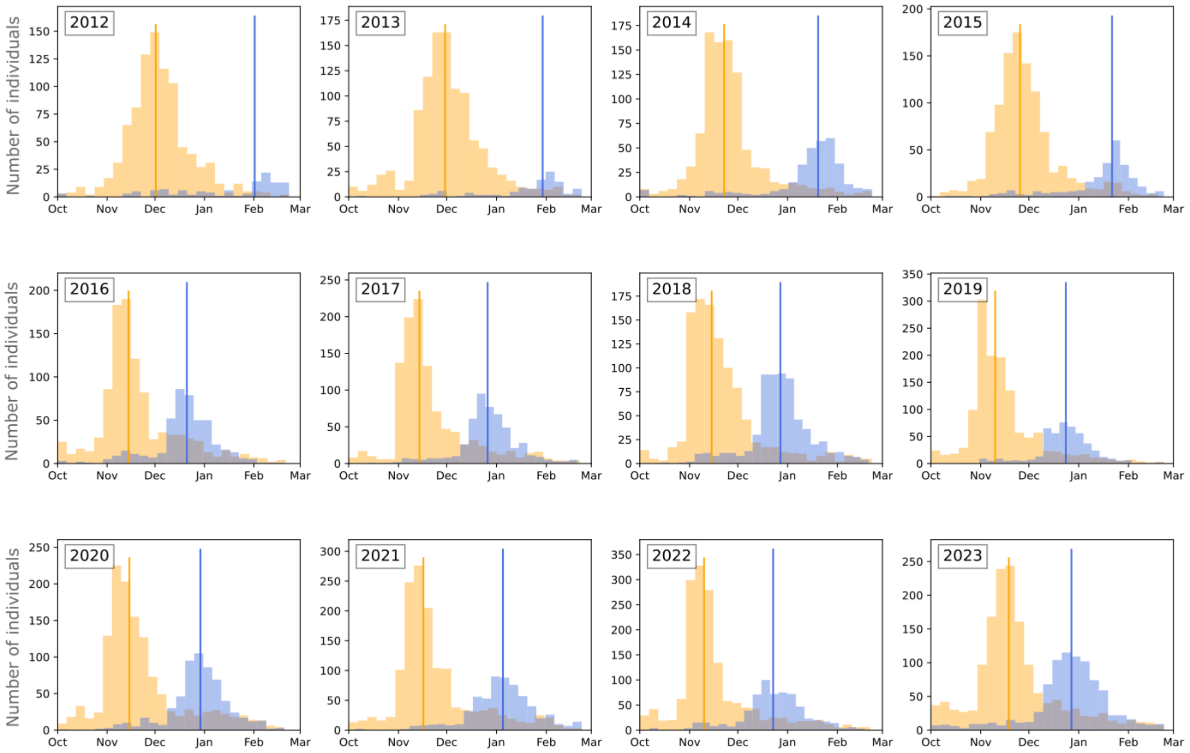


**Figure S1.** Breeding phenology in the King penguin colony of La Baie du Marin (Crozet Archipelago) in the past decade. Breeding dates were defined as the onset of the individual's first annual stay in the colony above 10 days after moulting, measured through RFID detection patterns (see Material and Methods for details). Blue bars represent the successful breeders from the previous year and orange bars represent both failed breeders from the previous year and individuals that didn't breed in the previous year. Vertical lines represent the median breeding date for the two groups.

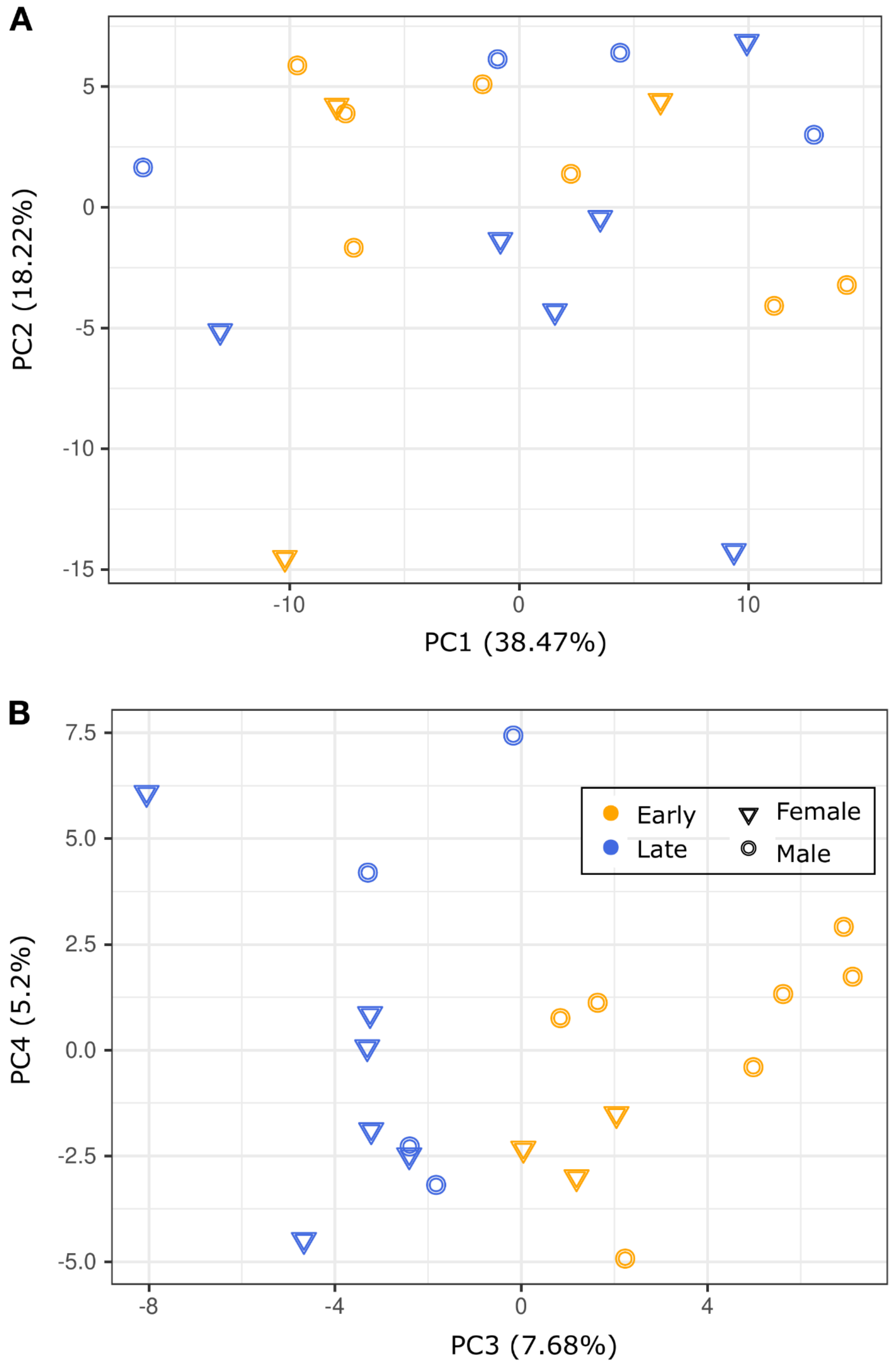


**Figure S2.** First four PC axes of the 500 most variable genes' expression of the 20 early- and late-hatchlings' blood RNA samples.

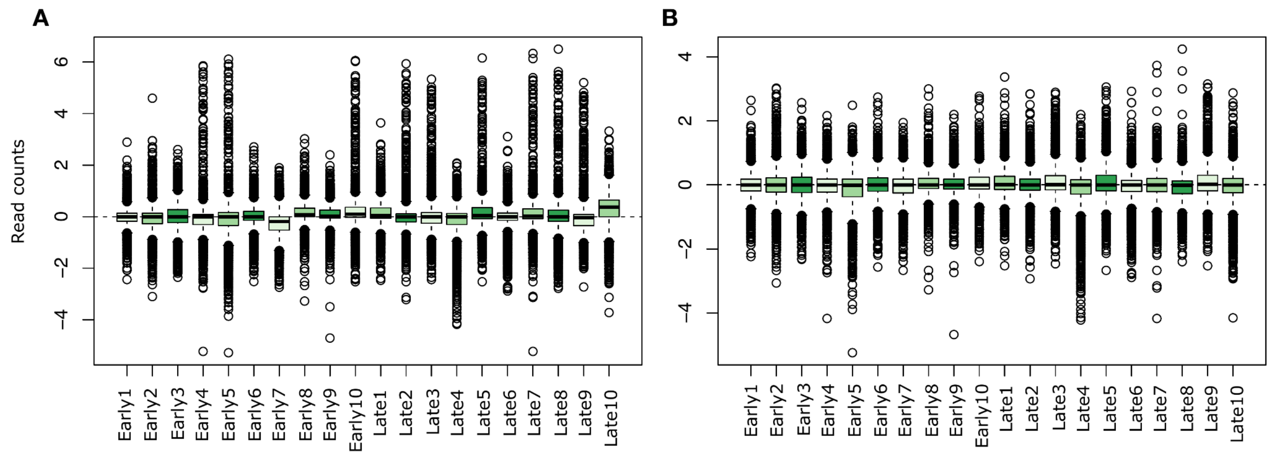


**Figure S3.** Gene expression counts per sample **A)** before and **B)** after normalisation with RUVSeq v1.38.0 (Risso et al. 2014).

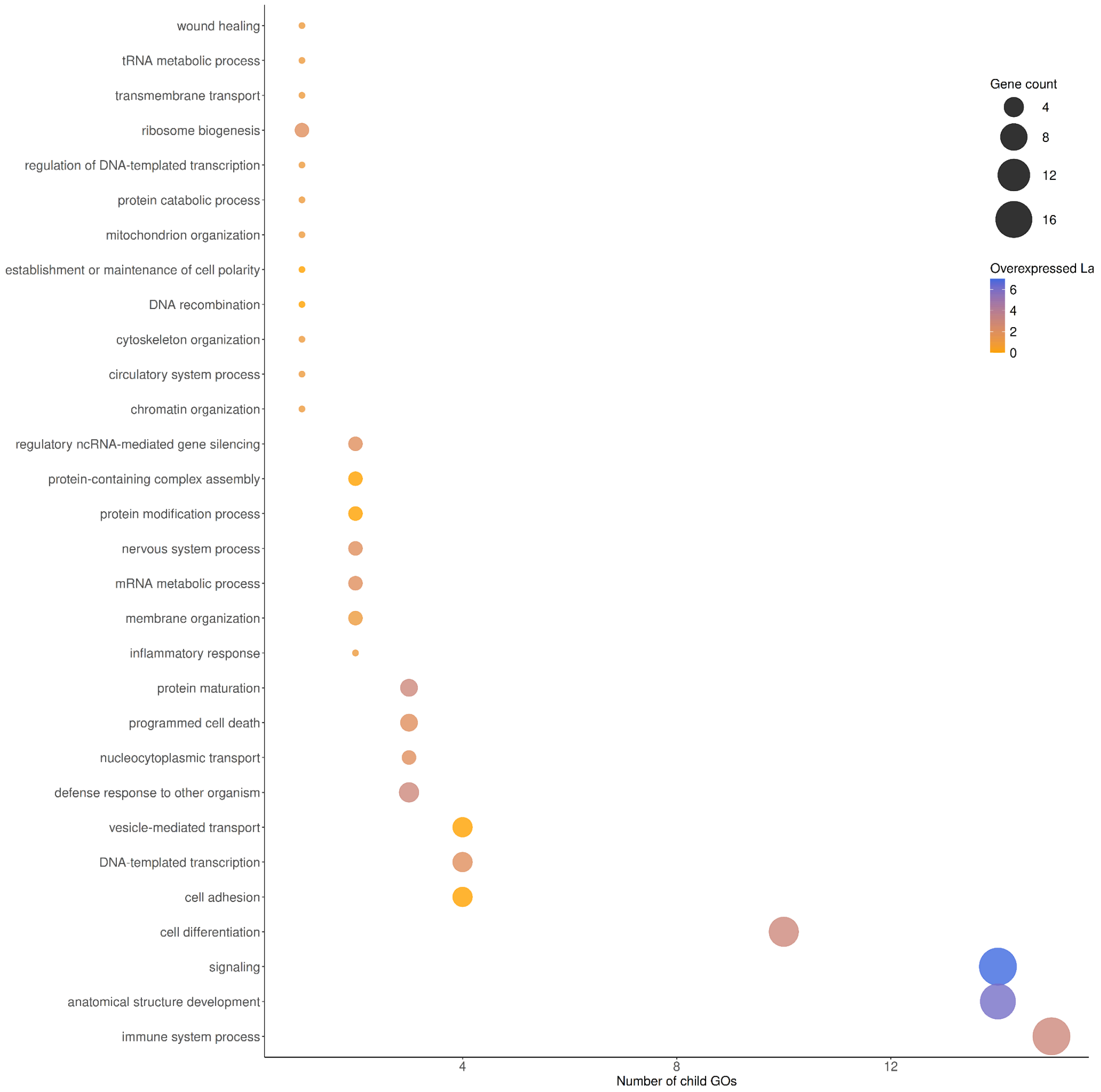


**Figure S4.** GO Slims enriched by the differentially expressed genes. Orange: overexpressed genes in early-born chicks; blue: overexpressed genes in late-born chicks. Intermediate colors: GOs enriched by both over- and underexpressed genes in late-born chicks.

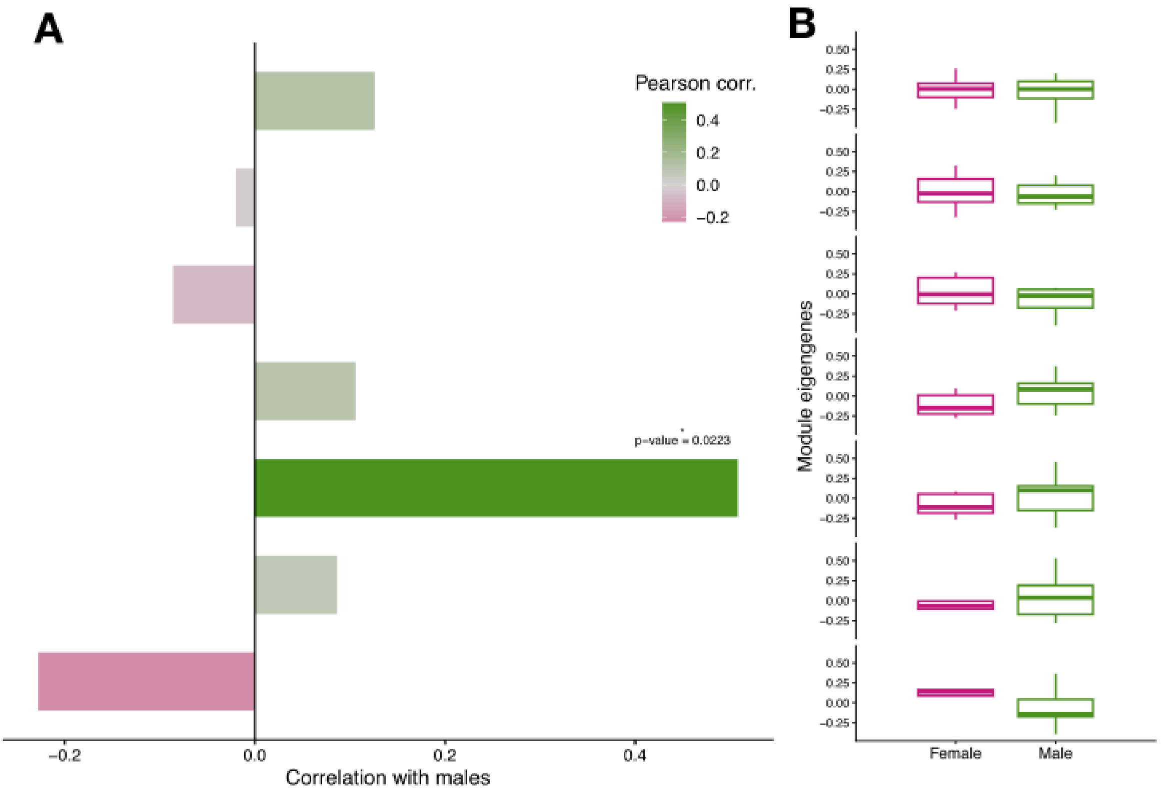


**Figure S5. Co-expression modules correlation with sex. A)** Pearson correlation between module eigengenes (ME) and sex (darker green and pink represents higher association with males and females, respectively); Detailed statistics in Table S6. **B)** Boxplots of female and male chicks' MEs of the normalised expression of genes belonging to each module (females: pink; males: green); Detailed statistics in Table S8.

### References


Bordier, Célia, Claire Saraux, Vincent A. Viblanc, et al. 2014. "Inter-Annual Variability of Fledgling Sex Ratio in King Penguins." *PloS One* 9 (12): e114052.

Dobson, F. S. 1992. "Body Mass, Structural Size, and Life-History Patterns of the Columbian Ground Squirrel." *The American Naturalist* 140 (1): 109–125.

Dutoit, Ludovic, Carina F. Mugal, Paulina Bolívar, et al. 2018. "Sex-Biased Gene Expression, Sexual Antagonism and Levels of Genetic Diversity in the Collared Flycatcher (Ficedula Albicollis) Genome." *Molecular Ecology* 27 (18): 3572–3581.

Fahlman, A., L. G. Halsey, P. J. Butler, et al. 2006. "Accounting for Body Condition Improves Allometric Estimates of Resting Metabolic Rates in Fasting King Penguins, Aptenodytes Patagonicus." *Polar Biology* 29 (7): 609–614.

Gunski, Ricardo José, Andrés Delgado Cañedo, Analía Del Valle Garnero, et al. 2017. "Multiple Sex Chromosome System in Penguins (Pygoscelis, Spheniscidae)." *Comparative Cytogenetics* 11 (3): 541–552.

Jakob, Elizabeth M., Samuel D. Marshall, and George W. Uetz. 1996. "Estimating Fitness: A Comparison of Body Condition Indices." *Oikos* 77 (1): 61–67.

Kriesell, Hannah J., Thierry Aubin, Víctor Planas-Bielsa, et al. 2018. "Sex Identification in King Penguins *Aptenodytes Patagonicus* through Morphological and Acoustic Cues." *The Ibis* 160 (4): 755–768.

Risso, Davide, John Ngai, Terence P. Speed, and Sandrine Dudoit. 2014. "Normalization of RNA-Seq Data Using Factor Analysis of Control Genes or Samples." *Nature Biotechnology* 32 (9): 896–902.

Sanchez, A. M., M. I. Flamini, K. Polak, et al. 2011. "Actin Cytoskeleton Remodelling by Sex Steroids in Neurones: Actin Cytoskeleton Remodelling by Sex Steroids in Neurones." *Journal of Neuroendocrinology* 24 (1): 195–201.

Saraux, Claire, Vincent A. Viblanc, Nicolas Hanuise, Yvon Le Maho, and Céline Le Bohec. 2011. "Effects of Individual Pre-Fledging Traits and Environmental Conditions on Return Patterns in Juvenile King Penguins." *PloS One* 6 (6): e20407.

Schulte-Hostedde, A. I., J. S. Millar, and G. J. Hickling. 2001. "Evaluating Body Condition in Small Mammals." *Canadian Journal of Zoology* 79 (6): 1021–1029.

Schulte-Hostedde, Albrecht I., Bertram Zinner, John S. Millar, and Graham J. Hickling. 2005. "Restitution of Mass–size Residuals: Validating Body Condition Indices." *Ecology* 86 (1): 155–163.
